# A *Drosophila* holidic diet optimised for growth and development

**DOI:** 10.1101/2024.09.24.614722

**Authors:** Sebastian Sorge, Victor Girard, Lena Lampe, Vanessa Tixier, Alexandra Weaver, Theresa Higgins, Alex P. Gould

## Abstract

Diets composed of chemically pure components (holidic diets) are useful for determining the metabolic roles of individual nutrients. For the model organism *Drosophila melanogaster,* existing holidic diets are unable to support rapid growth characteristic of the larval stage. Here, we use a nutrient co-optimisation strategy across more than 50 diet variants to design HoldFast, a holidic medium tailored to fast larval growth and development. We identify dietary amino acid ratios optimal for developmental speed but show that they compromise survival unless vitamins and sterols are co-optimised. Rapid development on HoldFast is not improved by adding fatty acids but it is dependent upon their *de novo* synthesis in the fat body via *fatty acid synthase* (*FASN*). HoldFast outperforms other holidic diets, supporting rates of growth and development close to those of yeast-based diets and, under germ-free conditions, identical. HoldFast provides new opportunities for studying growth and metabolism during *Drosophila* development.

## Introduction

Diet profoundly impacts physiology, metabolism and disease. Many decades of dietary research have had important consequences for public health including recommendations for the dietary intakes/allowances of most nutrients^1–4^. During the last century, classical studies identified nutrients that can only be obtained from the diet, such as essential amino acids and vitamins, versus those that the body can also synthesize^5,6^. The concentrations of individual essential and non-essential dietary nutrients as well as their relative amounts can influence metabolic and physiological parameters including lifespan^7–9^. Importantly, nutritional requirements are not fixed but change according to the specific pattern of metabolic demands at each stage of life. For example, fast growth rates during development place a particularly high demand on the quality of the diet. This is reflected in increased requirements for many essential micro- and macronutrients during pregnancy. Hence, some nutrients that are not categorized as essential during adulthood, such as arginine, become conditionally essential in the diet during fetal and postnatal development^5,10,11^.

Typical laboratory diets for most model organisms used in research are oligidic, containing crude natural ingredients that are complex and not chemically defined^12^. Most research using the fruit fly, *Drosophila melanogaster*, uses oligidic diets and these can vary considerably between laboratories (examples in ^13,14,15^). This lack of standardization, together with source/batch variations in the natural ingredients, make it challenging to compare results across different studies and contributes to reproducibility issues^16,17^. A common denominator in most oligidic diets is that the major protein source is an inactivated strain, or extract, of the yeast *S. cerevisiae*^18^. Yeast and other natural ingredients in the diet, such as cornmeal, contribute varying amounts of all three macronutrients (protein, carbohydrate and lipids) as well as essential micronutrients (vitamins and minerals). Oligidic diets containing hydrolysed yeast and sucrose/glucose have proved very useful for identifying the mechanisms by which dietary restriction (food dilution) extends lifespan^19–21^. Similar diets have also been combined with an analysis framework called nutritional geometry to demonstrate substantial differences in the protein-to-carbohydrate ratios that maximise longevity, fecundity and male attractiveness^22,23^. This illustrates the important point that diets optimised for one physiological parameter can be suboptimal for another.

Holidic diets composed entirely of chemically defined components offer great potential for *Drosophila* nutrition research^24^. The ability to manipulate independently the concentrations of each macro- and micronutrient molecule allows their specific metabolic and physiological functions to be clearly determined. The first near-chemically defined diet for *Drosophila* was described in 1946 and subsequently refined over many decades by numerous other studies^25–38^. These defined diets were instrumental in showing that micronutrients such as choline and the B-vitamins are essential but not Vitamin C or fat soluble vitamins^28,30,31^. They have also been used to demonstrate that dietary sterols but not dietary fatty acids are necessary for survival^28,33,39^. An important milestone in the development of complete holidic diets for *Drosophila* was the replacement of the protein source, casein, with individual amino acids^32^. This allowed investigation of the essentialities of amino acids as well as their optimal dietary balance^34–37,40^. An approach known as exome matching has calculated the ratios of dietary amino acids based on those obtained by *in silico* translation of the genome^36^. Exome matching to the *Drosophila* genome gave significant improvements over Hunt 1970 and other diets in terms of increasing reproductive fitness (fecundity) without a lifespan trade-off^32,36^. The latest holidic diets formulated with individual amino acids are impressive in their ability to support physiological traits of adult *Drosophila* to a similar degree as yeast-based oligidic diets^34–36^.

## Design

Existing holidic diets perform well for adult traits but are not widely used for pre-adult studies because they cannot support anywhere near the rate of development obtained with oligidic diets. The total duration of embryonic, larval and pupal development with an existing exome-matched holidic diet is at best ∼15 days, rather than the ∼10 days obtained with yeast-based diets^36^. As the durations of the non-feeding (embryonic and pupal) stages are relatively insensitive to dietary variation, this likely approximates to a halving of larval developmental speed and growth rate. Similar or slightly shorter delays in larval development also occur with other holidic diets, including the recently reported 108N pre-adult diet^28,33,35,38^. These substantial larval delays reflect compromised growth and metabolism, limiting the usefulness of holidic diets for *Drosophila* developmental biologists. Here, we systematically optimise an existing holidic diet^37^ for three key traits – mass gain (a measure of growth), larval duration (a measure of developmental speed), and the percentage of larvae completing development (a measure of survival). Via the systematic testing of more than 50 new diet formulations, successive improvements are made in the absolute and relative concentrations of macro- and micronutrients. Given the large number of formulations tested, the study design used multiple replicate vials for all dietary modifications but only those that yielded improvements were reproduced using multiple independent experiments done on different days. Using this strategy, we find that co-optimisation of individual amino acids, carbohydrates, lipids, vitamins and agarose is able to diminish substantially the delay associated with development on existing holidic diets. The culmination of this work is HoldFast, the first holidic diet to our knowledge that supports fast rates of *Drosophila* growth and development close to those of a yeast based diet.

## Results

### Amino acid optimisation accelerates development

We set out to identify the optimal balance of amino acids for larval growth and development on a holidic chemically defined diet (CDD). The starting point was CDD22, a diet we previously developed for adult *Drosophila*^37^ (**Supplemental Table 1**). All of the ingredients in CDD22 are high-purity chemicals, including the gelling agent 2-hydroxyethyl agarose. This is a chemically defined replacement for the agar used in many holidic diets, which not only contains agarose but also agropectin and several impurities. First, three separable developmental parameters (speed, growth, and survival) were determined at 25°C for larvae of a control strain, *Wolbachia* free *w^11^*^18^ *iso31* raised on CDD22 and benchmarked against a standard diet (STD) of yeast-glucose-cornmeal-agar^13^. We observed that the time taken for 50% of surviving larvae to develop into puparia, time-to-pupariation 50 (TTP_50_), was ∼3.3 days longer on CDD22 (7.63 ± 0.2 days) than on STD (4.3 ± 0.18 days), equating to a 77% slower pace of larval development (**Figures 1A** and **1B**). CDD22 not only has a longer TTP_50_ but also a reproducible small deficit in the percentage of larvae surviving to pupariation (CDD22: 92% ± 2.2%, STD: 96.2% ± 1.3%, **Figure 1C**). CDD22 also supports substantially less growth (mass gain) during overall development than the yeast-based diet, with 2-4 day old adult females and males weighing an average of 0.71 mg and 0.56 mg respectively, compared to 1.43 mg and 0.89 mg for those raised on STD (**Figure 1C**). We conclude that CDD22 supports larval developmental speed, growth and survival much less well than a yeast-based diet. In terms of speed and survival, CDD22 does perform slightly better but roughly within the same range as other published holidic diets^34–36^. These results demonstrate that considerable optimisation of holidic diets is required if they are to support anywhere near the rates of larval development obtained with yeast-based diets.

**Figure 1.**
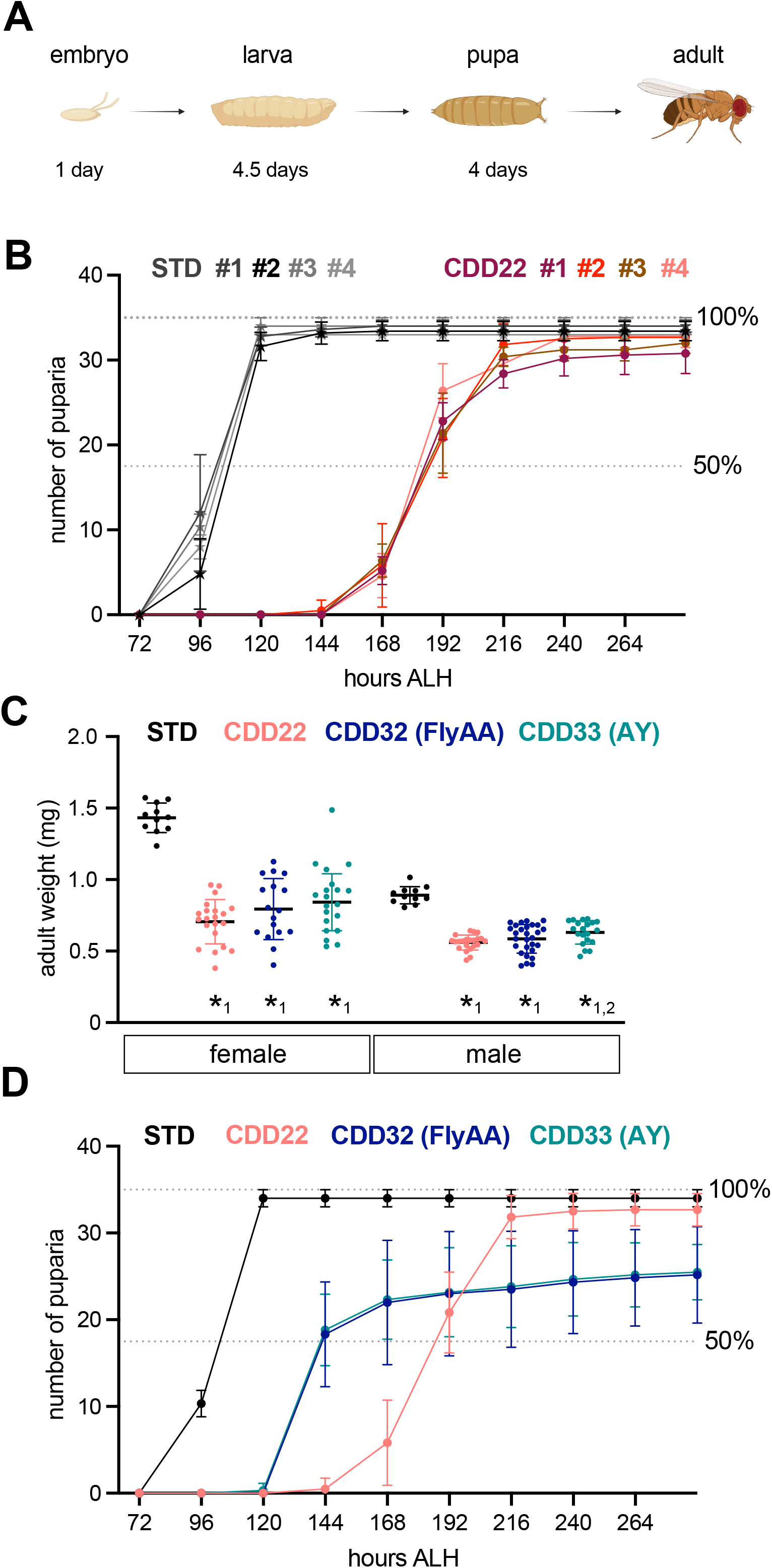
Larval developmental speed, growth and survival on STD versus CDD22-based diets. (A) Schematic of *Drosophila* development with approximate durations of each developmental stage on an optimal undefined yeast-based diet (such as STD) at 25 °C. (B) Time-to-Pupariation (TTP) curves of *w^1118^ iso31* larvae raised on standard fly food (STD) or the initial chemically defined diet CDD22. Data shown are from four independent experiments (#1 to #4). (C) Adult male and female weights at 2 to 4 days after eclosion for *w^1118^ iso31* flies raised as larvae on STD, CDD22 and CDD22-base diets with the amino acid profiles of FlyAA (CDD32) or autolysed yeast (AY, CDD33). The weights of adult males and females raised on all holidic diets are significantly less than those raised on STD (*1, p<0.001). Males raised on CDD33 weigh significantly less than those raised on CDD22 (*2, p=0.0133). (D) Time-to-Pupariation (TTP) of *w^1118^ iso31* larvae raised on STD, CDD22 or CDD22-based diets with the amino acid profiles of FlyAA (CDD32) or AY (CDD33).

To identify potential differences that could underlie the different larval performances of CDD22 and STD, we compared their macronutrient compositions. STD diet has 58.5 g glucose per litre and chemical analysis indicates that it also contains 10.5 g protein per litre, with roughly 2/3 of the protein coming from autolysed yeast (AY, *S. cerevisiae*, containing 31.25 g total AA per 100g) and 1/3 from cornmeal (containing 6.16 g total AA per 100g, **Supplemental Table 2**). In comparison, CDD22 is formulated with 10 g sucrose and 25 g of total amino acids per litre. The very different macronutrient contents of the two diets equate to protein:carbohydrate mass ratios of <0.18 for STD and 2.5 for CDD22. Both the total amount and the relative proportions of dietary amino acids are key determinants of the rates of larval growth and development^36,41–44^. Given that the total concentration of amino acids in CDD22 is more than double that of STD, this parameter alone cannot be rate-limiting for development on the holidic diet. To determine the relative proportions of individual amino acids in STD and its main components cornmeal and autolysed yeast (AY), chemical hydrolysis was used and this revealed an amino acid profile very different from CDD22 yet surprisingly similar to the *in silico* translated *Drosophila* exome used in the FlyAA holidic diet^36^ (**Figure S1A**). In addition, the amino acid profiles of the yeast proteome and the *in silico* translated yeast exome are also quite similar (**Figure S1B**). Consistent with this, the 25 most highly expressed proteins in the yeast genome have an amino acid profile that is not very different from that of the translated exome (**Figure S1B**). This is reflected in Spearmann’s rank correlation coefficients of 0.86 and 0.89 for comparisons of the exome with the proteome or with the top 25 proteins respectively (**Figure S1C**). We conclude that the amino acid profiles of the *in silico* translated genome and the proteome are highly correlated in yeast and likely also in other species. Two new versions of CDD22, termed CDD32 and CDD33, were therefore designed keeping the total mass of amino acids constant but adjusting their balance to that of FlyAA or AY respectively (**Supplemental Table 1**). However, neither CDD32 nor CDD33 improved the developmental growth deficit of CDD22 compared to STD diet and they were ∼20% worse than CDD22 in terms of survival to pupariation (**Figures 1C** and **1D**). Despite this, both CDD32 and CDD33 accelerated larval development by an impressive and almost identical amount of ∼2 days compared to CDD22 (**Figure 1D**). Hence, matching the amino acid profile of the CDD22 base diet to either autolysed yeast or to the *in silico* translated *Drosophila* exome dramatically improves developmental speed but not growth or survival.

The unexpectedly similar amino acid ratios and dietary performances of autolysed yeast and the fly exome prompted us to make more extensive amino acid variations. We therefore formulated additional holidic diets based upon CDD22 but containing amino acid profiles previously reported as mismatched^36^, CDD-MM1 and CDD-MM2, or containing an amino acid profile intermediate between that of the abundant hemolymph protein, larval serum protein 1 beta (Lsp-1β), and autolysed yeast (CDD-MM3, **Supplemental Table 1**). Surprisingly, CDD-MM1 and CDD-MM3 outperformed CDD22 in terms of TTP whereas CDD-MM2 was considerably worse and also had poor survival (**Figure S2A**). To clarify which amino acid ratios correlate with fast development, we used principle component analysis (PCA) to assess variation between high and low performance diets versus the translated exomes of multiple species. This analysis included the amino acid compositions of 8 different diets or their components (5 holidic diets, STD, cornmeal and autolysed yeast) together with those of the translated exomes of 18 species spanning all domains of life, including many model organisms. PC1 (29.5% of variance) separated low performance diets (CDD22 and CDD-MM2) away from a large cluster of high performance diets and translated exomes (**Figure S2B**). It did not, however, separate species-specific differences between translated exomes. The loadings plot indicates that Glu, Val, Ile and Ser make large contributions to PC1 separation (**Figure S2C**). PC2 (22.5% variance) does separate species-specific translated exomes, as well as high performance diets, and it is largely driven by Tyr, Phe, Lys and Met (**Figures S2B** and **S2C**). Together, the experimental and PCA findings demonstrate that diets that cluster with the majority of translated exomes along PC1, including the “mismatched” diets CDD-MM1 and CDD-MM3, tend to support fast larval development. In contrast, those diets that separate away from translated exomes along PC1, such as CDD22 and CDD-MM2, are only capable of supporting slow development.

### Vitamin optimization increases survival

The findings thus far show that amino acid optimization of the CDD22 base diet can speed up development but this comes at the expense of decreased survival and no improvement in overall growth. It therefore follows that other components of the CDD32/33 holidic diets, such as micronutrients (CDD22 micronutrients: vitamins, sterol, minerals and nucleotides, will be called µ1 hereafter), may be limiting for growth and survival at a fast pace of development. As STD supports optimal growth, development and survival, its active vitamin content and that of its individual complex ingredients (cornmeal and yeast) were analysed by microbiological bioassays (**Figure S3A**). Yeast not cornmeal makes the largest contributions (ranging from 76% to 98%) to all the vitamin bioactivity in STD for vitamin B1 (thiamine), vitamin B2 (riboflavin), vitamin B3 (niacin), vitamin B5 (pantothenate), vitamin B6 (pyridoxin), vitamin B7 (biotin), vitamin B9 (folate) and choline (**Figures S3A** and **S3B**). This analysis also revealed that the processing/cooking of STD strongly depletes (>60%) the bioactivities of vitamin B1 and B2 and moderately lowers (<54%) the bioactivities of vitamin B5 and B6 (**Figure S3C**). In contrast, food processing increased vitamin B7 and choline bioavailability in STD by ∼50%, potentially due to release from protein-bound or phospholipid moieties respectively^45,46^ (**Figure S3C**).

For all vitamins, the calculated concentrations in CDD22-based diets are greater than those measured by bioassay in STD but, in relative terms, vitamin B3 and choline are under-represented (**Figure S3D**). However, increasing vitamin B3 (niacin) or choline in the context of CDD33 by two- or five-fold, either individually or together, did not substantially improve survival or developmental speed, indicating that they are not limiting (**Figure S3E**). Surprisingly, swapping the original CDD22 vitamin mix (µ1) with that of the FlyAA diet^36^ increased TTP_50_ by ∼70 h, indicative of a long developmental delay (**Figure S3F**). As an alternative optimisation approach, a physiological multivitamin mix based on STD diet concentrations was established. This differs considerably from the original CDD22 vitamin mix, with substantially lower concentrations of vitamins B1, B2, B5, B6, B7 and B9. Larvae were then raised on reformulated versions of CDD33, with varying amounts of the physiological multivitamin mix ranging from 1x to 16x the concentrations measured in STD food. With 1x physiological vitamins, larvae did not survive on CDD33 and, even with 2x vitamins, less than 50% of larvae were able to complete pupariation (**Figure 2A**). However, 4x to 8x physiological multivitamin mix supported progressively increasing developmental speed and survival to pupariation (**Figure 2A**). The 8x physiological vitamin mix was able to increase survival on CDD33 from 71% to 82% while maintaining the same fast pace of development with a TTP_50_ of ∼150 hr (**Figure 2A**). Further increasing the vitamin concentration in CDD33 to 16x the STD value did not substantially improve survival but marginally increased developmental speed (**Figure 2A**). Hence, low levels of physiological vitamin mix in CDD33 compromise developmental speed but higher levels show an improvement in larval survival. These co-optimisation experiments reveal that micro- and macronutrient requirements are interdependent, with striking differences in the vitamin requirements of STD, FlyAA and CDD33 diets.

**Figure 2.**
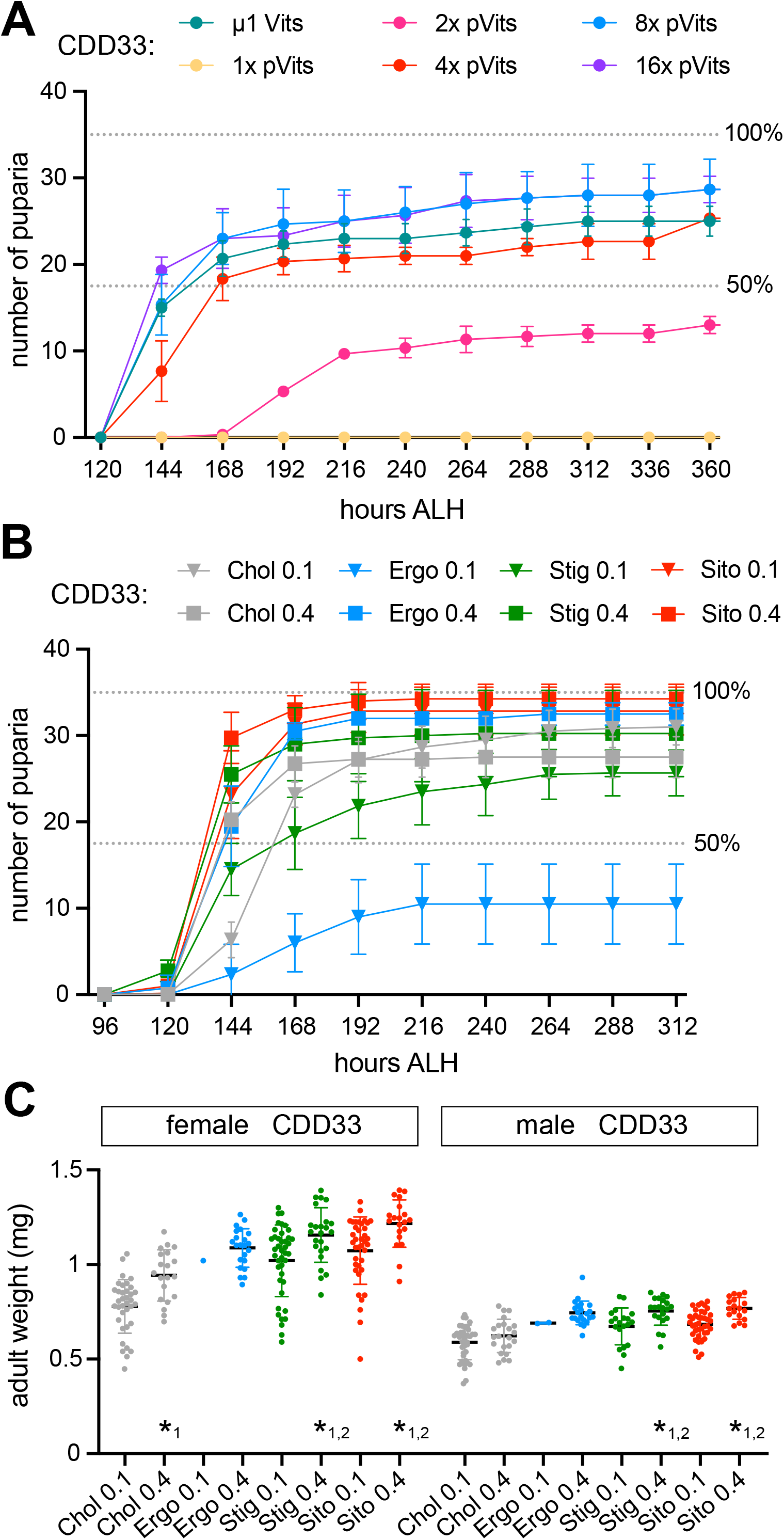
Vitamin and sterol optimisations increase survival, growth and development on a holidic diet. A) Time-to-Pupariation (TTP) of control (*w^1118^ iso 31*) larvae on CDD33 containing the original vitamin mix (μ1) or varying amounts (1x to 16x) of a physiological vitamin mix based on STD. Note that 1x refers to the bioactive amounts measured in STD, which are insufficient to support larval survival and development on CDD33. B) Time-to-Pupariation (TTP) of control larvae on CDD33 containing one of four different sterols: cholesterol (Chol), ergosterol (Ergo), stigmasterol (Stig) or beta-sitosterol (Sito) at a concentration of 0.1 g or 0.4 g per litre. C) Adult male and female weights at 2 to 4 days of eclosion from the sterol experiments in panel B. Large differences in weight are observed between different sterols and amounts, with 0.4 g per litre of beta-sitosterol and 0.1 g per litre of cholesterol giving the highest and lowest weights respectively. Statistical significance is indicated for pairwise comparisons between low and high doses of sterol (*1, p<0.01) and between high doses of cholesterol and another sterol (*2, p<0.001). Ergosterol was not statistically analysed due to the small number of surviving males.

### Sterol optimization increases developmental growth and survival

Insects like *Drosophila* are sterol auxotrophs, requiring dietary sterols for the biosynthesis of the ecdysteroid hormones essential for organismal growth and developmental transitions^47,48^. Current holidic diets, including the CDD22 base diet, contain a single animal-derived sterol, cholesterol. Nevertheless, different sterol species have different larval tissue distributions and are metabolised into distinct ecdysteroid variants that do not all support developmental progression equally well^48,49^. We therefore compared TTPs for four different sterols: the two common phytosterols (stigmasterol and β-sitosterol), the main sterol of yeast (ergosterol) and the zoosterol cholesterol. Two sterol concentrations were used, half the original amount in CDD33 (0.1 g/litre) or a four-fold increase (0.4 g/litre) which is at/above maximum solubility. Unlike amino acid optimisations, changing sterols did not substantially alter the time at which the first larvae started to pupariate, but increased sterol concentrations steepened the pupariation curve and decreased TTP_50_, indicative of greater developmental synchronisation (**Figure 2B**). Higher sterol concentrations also increased adult weight, demonstrating a major improvement in developmental growth (**Figure 2C**). Interestingly, we also observed large differences between the four different sterols in terms of adult weight and particularly larval survival (**Figures 2B** and **2C**). Surprisingly, at 0.1 g/litre, the major sterol in yeast (ergosterol) supported survival and pupariation synchrony far less well than the three other sterols. Nevertheless, the few adults that did eclose on CDD33 with 0.1g/l ergosterol were not substantially lighter, suggesting that the primary developmental deficit may be survival not growth (**Figure 2C**). Among the four sterols, β-sitosterol showed the best performance, with 0.4 g/litre supporting near 100% survival, a TTP_50_ of ∼130 hr and overall developmental growth of 0.77 mg ± 0.06 mg and 1.22 mg ± 0.13 mg for males and females respectively (**Figures 2B** and **2C**). Nevertheless, a combination of 0.15 g/litre of cholesterol and β-sitosterol gave a similar TTP_50_ as 0.3 g/litre of β-sitosterol (**Figure S3G**). For practical reasons (see Discussion), we therefore selected 0.2 g/litre of β-sitosterol plus 0.2 g/litre cholesterol as our optimal sterol mix for optimising developmental growth, speed and survival on CDD33 based diets. This improved sterol mix was then combined with the physiological 8x vitamin mix to give an optimised micronutrient mix (µ2), which we used to create CDD40 (autolysed yeast amino acid ratios with µ2 micronutrients). We also generated CDD41, a variant of CDD40 with the same total amino acid content but lower concentrations of five amino acids (Asn, Gln, Gly, Pro and Thr) that are substantially higher in the hemolymph of larvae raised on CDD40 than those raised on STD (**Figure S1D** and **Supplemental Table 1**). The amino acid profile of CDD41 is also slightly more similar to that of most translated exomes than to CDD40 (**Figure S2B**).

To further investigate the interdependence of macro- and micronutrients on development, we tested the combined effects of optimised sterols and vitamins on three of the amino acid mismatched diets that perform poorly: CDD22, CDD-MM1 and CDD-MM2 (**Figure S2A**). Adult weight measurements indicated that switching from the initial micronutrient mix (µ1) to the optimised version (µ2) was able to improve the overall growth during development of females on all three mismatched diets (**Figure S3H**). Optimised micronutrients did not, however, improve TTP_50_ or survival on CDD22 and, for CDD-MM2, both parameters were substantially worse (**Figure S3I**). In contrast, for CDD-MM1, the optimised micronutrient mix improved survival, shortened TTP_50_ and increased male and female weight (**Figures S3H** and **S3I**). Together, the dietary vitamin and sterol manipulations highlight that micronutrient optimisation has a general and major growth-promoting effect independent of amino acid ratios, but that effects exerted by micronutrients on developmental speed are dependent on the amino acid ratio.

### Dietary carbohydrate promotes developmental growth and speed but not survival

STD has a higher carbohydrate content than all CDDs and this could contribute to its high performance. STD contains the starch in cornmeal and glucose at 58.5 g/litre, whereas CDD diets contain 10 g/litre of sucrose and no other carbohydrate. Consistent with the non-essential nature of dietary carbohydrates, varying the sucrose content of CDD40 between 0 and 40 g/l did not significantly alter larval survival (**Figure 3A**). In contrast, this dietary manipulation did markedly alter both the TTP_50_ and overall growth during development (**Figures 3A** and **3B**). A sucrose concentration of 10-20 g/litre was optimal for TTP_50_ and 20 g/litre was optimal for overall male growth (**Figures 3A** and **3B**). Interestingly, sucrose concentrations lower or higher than 10-20 g/litre both slowed development (**Figure 3A**). This defines 20 g sucrose per litre as the optimal dietary concentration, which we later used in CDD48 (see below). Starch supplementation of CDD41 did not, however, significantly alter TTP_50_ or survival of *w^11^*^18^ *iso31* larvae (**Figure S3A**).

**Figure 3:**
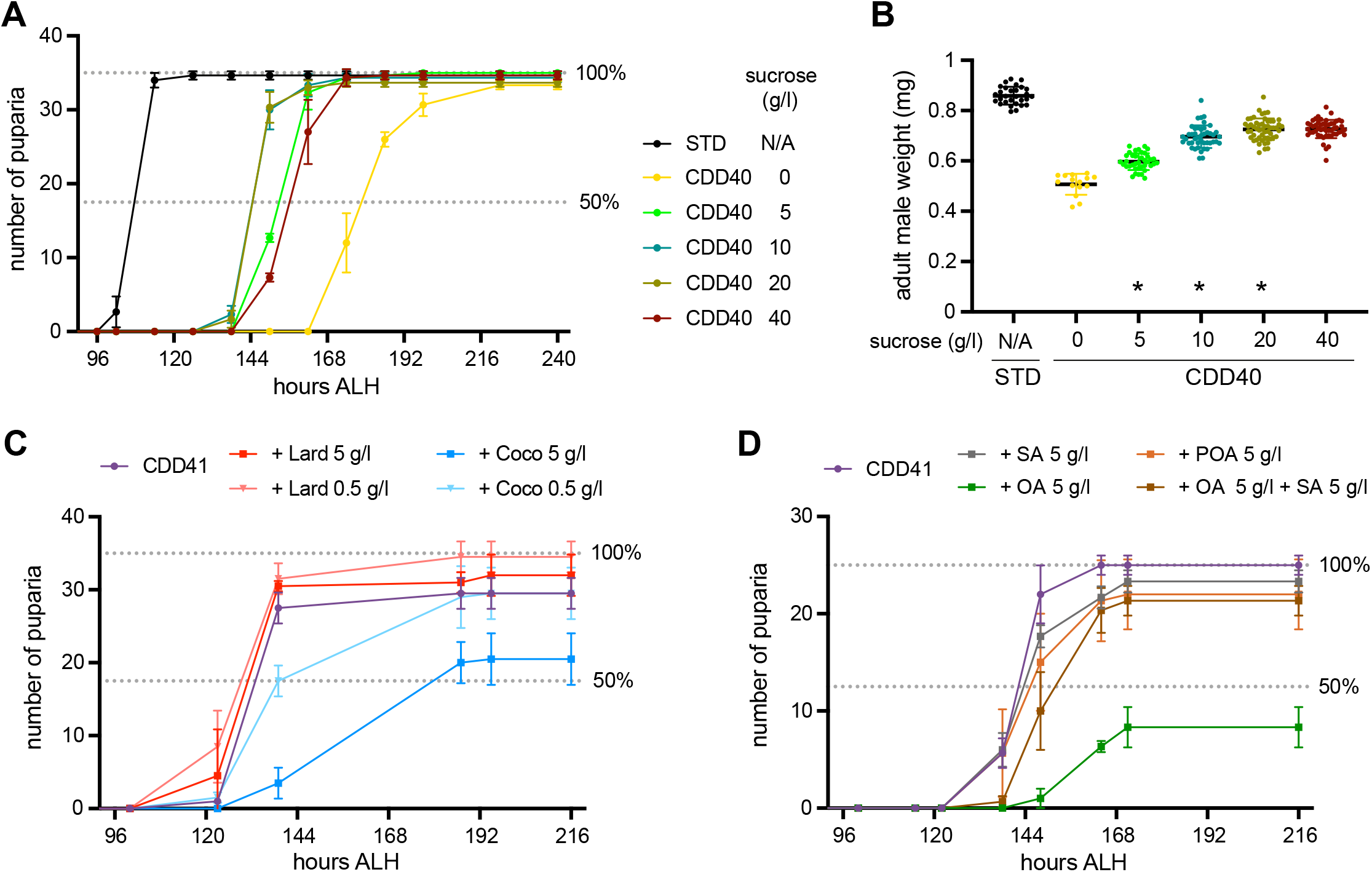
Carbohydrate optimisation and effects of fats on CDD performance. A) Time-to-Pupariation (TTP) of wild-type (*Oregon R)* larvae on STD or CDD40 supplemented with the indicated sucrose concentrations. TTP_50_ was shortened by sucrose concentrations up to 20 g/l but lengthened by a further increase to 40 g/l. B) Weights of individual male adult flies from the TTP experiment in Fig 3A. Weight increases with increasing sucrose, plateauing between 20-40 g/litre. Statistical significance is indicated for pairwise comparisons between each sucrose concentration and the next higher one (*, p<0.01). C) TTP of control (*w^1118^ iso31*) larvae on CDD41 supplemented with coconut oil (Coco) or lard. Lard at 0.5 g/l moderately increased larval survival whereas both 0.5 g/l and 5 g/l of coconut oil delayed development and decreased survival. D) TTP of *Oregon R* (*OreR*) larvae on CDD41 supplemented with 5 g/l of the indicated free fatty acids (stearic acid, SA; oleic acid, OA; palmitoleic acid, POA). Monounsaturated OA but not monounsaturated POA delays development and decreases survival, which is largely rescued by the presence of saturated SA.

### Larvae raised on HoldFast are sensitive to dietary lipid imbalances

CDD40 and all subsequent CDD formulations in this study contain optimized sterols but, like other holidic diets, they are free from all non-sterol lipids including fatty acids. Dietary fatty acids are, however, abundant in STD and so we tested whether their addition to CDD41 might improve its performance. Surprisingly, however, supplementation with a natural source of triacylglycerols, coconut oil, delayed development and decreased survival although the addition of lard did give a minor increase in survival (**Figure 3C**). Importantly, neither fat source substantially altered overall growth during development, as determined by adult male weights (**Figure S3B**). Additional lipid supplementation experiments with each of the major 18 carbon (C18) fatty acids in lard revealed that a low concentration (0.5 g/l) of stearic (C18:0), oleic (C18:1) or linoleic (C18:2) acid did not substantially alter TTP_50_, survival or overall male growth (**Figures S4B** and **S4C**). However, a ten-fold higher concentration (5 g/l) of oleic or linoleic acid was incompatible with larval survival (**Figure S4C**). In contrast, stearic acid, a saturated fatty acid, was well tolerated and did not significantly alter developmental speed or overall growth (**Figures 3D**, **S4B** and **S4C**). We also found that high concentrations of oleic or linoleic acid were considerably more detrimental for development on a holidic than on a STD diet (**Figure S4D,** compare with **S4C**). Interestingly, on CDD41, the larval lethality of oleic acid was largely rescued by additional supplementation with an equimolar amount of stearic acid (**Figure 3D**). Hence, a dietary lipid imbalance with a high ratio of unsaturated to saturated fatty acids can impact negatively on larval development, especially on holidic diets. Given that dietary fatty acids gave no major detectable benefits for development, they were omitted from the optimised holidic diet.

### Lipid synthesis in the fat body is essential for development on holidic but not STD diet

Metabolic networks are highly flexible and adapt to dietary variations in order to maintain biochemical and energetic homeostasis^50–52^. It is therefore inevitable that larval metabolism will differ significantly between holidic and STD diets. For CDD41 and related diets to be useful for nutritional studies, it is particularly important to define how metabolism adapts to the complete absence of dietary fatty acids. A key question here is how holidic diets change the genetic requirements for fatty acid synthases, enzymes required for the *de novo* synthesis of fatty acids from dietary carbohydrates and amino acids. *Drosophila* has three annotated fatty acid synthases including *fatty acid synthase 1* (*FASN1*), an essential gene expressed in many tissues^53^ and *FASN2*, involved in synthesis of branched-chain fatty acids^54^. Midgut enterocytes are an important site for dietary lipid processing in larvae^55–57^ but enterocyte-specific knockdown of *FASN1* and *FASN2* (*NP1>FASN[RNAi]*) was fully viable to adulthood on both STD and CDD41 (**Figures 4A** and **4B**). *Drosophila* adipose tissue (fat body) is the major site where fatty acids are synthesized and stored as triacylglycerols in lipid droplets^58,59^. Nevertheless, *FASN1* is not required in the fat body for larval survival on a yeast-based oligidic diet^53^. Consistent with this, fat body-specific knockdown of FASN1+2 (*Lpp>FASN[RNAi]*) permitted the majority (80%) of larvae raised on STD to survive to pupariation, although there was ∼80% lethality at/around adult eclosion (**Figures 4A** and **4B**). In contrast, when these knockdown larvae were raised on CDD41 only ∼30% survived to pupariation and none eclosed as adults (**Figures 4A** and **4B**).

**Figure 4:**
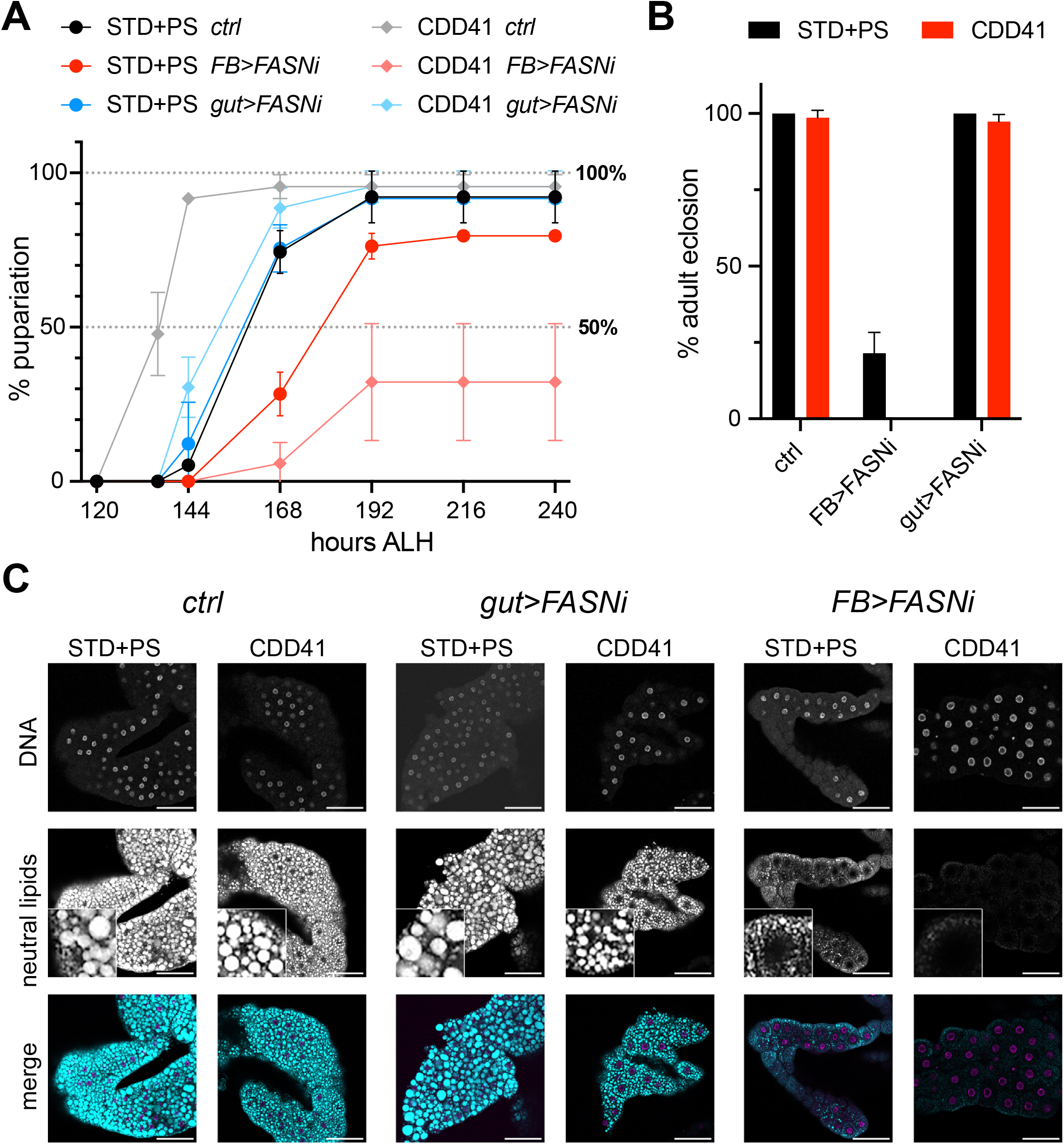
Fat body *FASN* is essential for the completion of development on CDD but not STD. A) Time-to-Pupariation (TTP) on STD or CDD41, both with penicillin and streptomycin (STD+PS or CDD41 respectively). Larvae are control (*ctrl*) genotype (*Myo1A-Gal4* and *apoLpp-Gal4*), enterocyte-specific FASN1+2 knockdown (*gut>FASNi* using *Myo1A-GAL4*), or fat body-specific FASN1+2 knockdown (*FB>FASNi* using *apoLpp-GAL4*). For fat-body FASN1+2 knockdown larvae, the average percentage pupariations and standard deviations are 80 % ± 0.6 % on STD+PS and 32% ± 19 % on CDD41. B) Adult survival (% adult eclosion) for control genotype (*ctrl*), with FASN1+2 knockdown in gut enterocytes (*gut>FASNi*), or in the fat body (*FB>FASNi*). The adult-lethal phenotype of fat-body FASN knockdown is partially penetrant on STD+PS but completely penetrant on CDD41(+PS). C) Confocal projections of third-instar larval fat bodies stained for neutral lipids (using LipidTOX dye) and nuclei/DNA (with Hoechst 33342 dye). Lipid droplets tend to be smaller in control (*NP1>, Lpp>*) larvae raised on CDD41(+PS) than on STD+PS. Fat body-specific (*FB>FASNi*) but not enterocyte-specific (*gut>FASNi*) FASN1+2 knockdown depletes LDs more strongly on CDD41(+PS) than on STD+PS. Inserts in LipidTOX channel are a 4x magnification. All scale bars are 100 μm.

We next tested whether lipids present in STD are responsible for rescuing the lethality of fat body-specific FASN1+2 knockdown. The major contribution of fatty acids to STD diet comes from the cornmeal. We therefore supplemented holidic diet with corn oil and observed that this decreased survival of control larvae and adults (**Figures S5A** and **S5B**). Similar detrimental effects were previously observed with oleic and linoleic acid supplementation of CDD41 (**Figures 3D** and **S4C**). However, corn oil supplementation almost fully rescued the development and adult survival of FASN1+2 knockdown larvae (**Figures S5A** and **S5B**). Similarly efficient rescue was also observed with palmitic acid (C16:0), the major product of the FASN reaction but, interestingly, not with a longer-chain fatty acid: stearic acid (C18:0) (**Figures S5C** and **S5D**). Hence, fat body *FASN1+2* are only essential for development in the absence of dietary fatty acids. We noticed that fat body-specific knockdown of FASN1+2 decreased the opacity of the fat body and, to investigate further, it was stained with a neutral lipid dye (LipidTOX). Both the large perinuclear and small cortical lipid droplets in the fat body^60,61^ are labelled with this dye in control genotype larvae raised on STD (**Figure 4C**). When control larvae are raised on CDD41, both categories of lipid droplets appear to be moderately smaller, perhaps reflecting decreased triacylglycerol storage by the fat body (**Figure 4C**). Gut enterocyte-specific knockdown of FASN1+2 (*Myo1A>FASN[RNAi]*) did not noticeably alter the size or abundance of lipid droplets in the fat body of larvae raised on STD or the holidic diet (**Figure 4C**). In contrast, fat body-specific knockdown of FASN1+2 (*apoLpp>FASN[RNAi]*) had a dramatic effect on lipid droplets. For larvae raised on STD, large perinuclear lipid droplets were depleted and for those raised on CDD41, almost all droplets were missing (**Figure 4C**). These striking findings together demonstrate that fatty acid synthesis within the larval fat body is strictly required for survival and neutral lipid accumulation on CDD41 but not on STD.

### Microbiota accelerate development on STD but not CDD

CDD40 and CDD41 are substantially improved over CDD22 in terms of larval development but we wanted to investigate why their TTP_50_ still remains ∼30 hr longer than for STD. Two antibiotics, penicillin and streptomycin (PenStrep), were used in all formulations of CDD22 to CDD41 because they are necessary to prevent opportunistic bacterial infection of holidic diets. However, we reasoned that these antibiotics must also interfere with the endogenous *Drosophila* microbiome, which can itself accelerate larval development^62–66^. STD diet does not contain PenStrep but, like CDD formulations, it does contain a broad-spectrum antimicrobial (nipagin/methylparaben) and an antifungal (bavistin/carbendazim). Nevertheless, flies raised on STD maintain a functional microbiota containing species representing the two major genera: *Acetobacter* and *Lactiplantibacillus*^67^.

To test whether microbiota could account for the slower speed of holidic versus yeast-based diets, we first supplemented STD with PenStrep at either the same (0.25 g/litre) as used in the CDD formulations or a ten-fold lower (0.025 g/litre) concentration. Both PenStrep concentrations slowed the TTP_50_ of *w*^*1118*^ *iso 31* control larvae on STD from ∼102 hr to ∼132 hr, which is close to the value obtained with CDD40 (**Figure 5A**). PenStrep addition to STD also produced a comparable lengthening of the TTP_50_ of larvae of the *white^+^* wildtype strain, *Oregon R* (**Figure S6A**). As an alternative to antibiotics, the effect of germ-free conditions was tested via sterile transfer of *w*^*1118*^ *iso 31* embryos to STD vials, and this similarly lengthened the TTP_50_ close to that of CDD40 (**Figure 5B**). Furthermore, under these sterile transfer conditions, PenStrep has little or no effect on TTP_50_ on STD or CDD40, indicating that the antibiotic delay is due to microbial inhibition not some other toxic side-effect (**Figure 5B**). Together, these results show that the microbiome is required for development to proceed faster on STD than on CDD.

**Figure 5:**
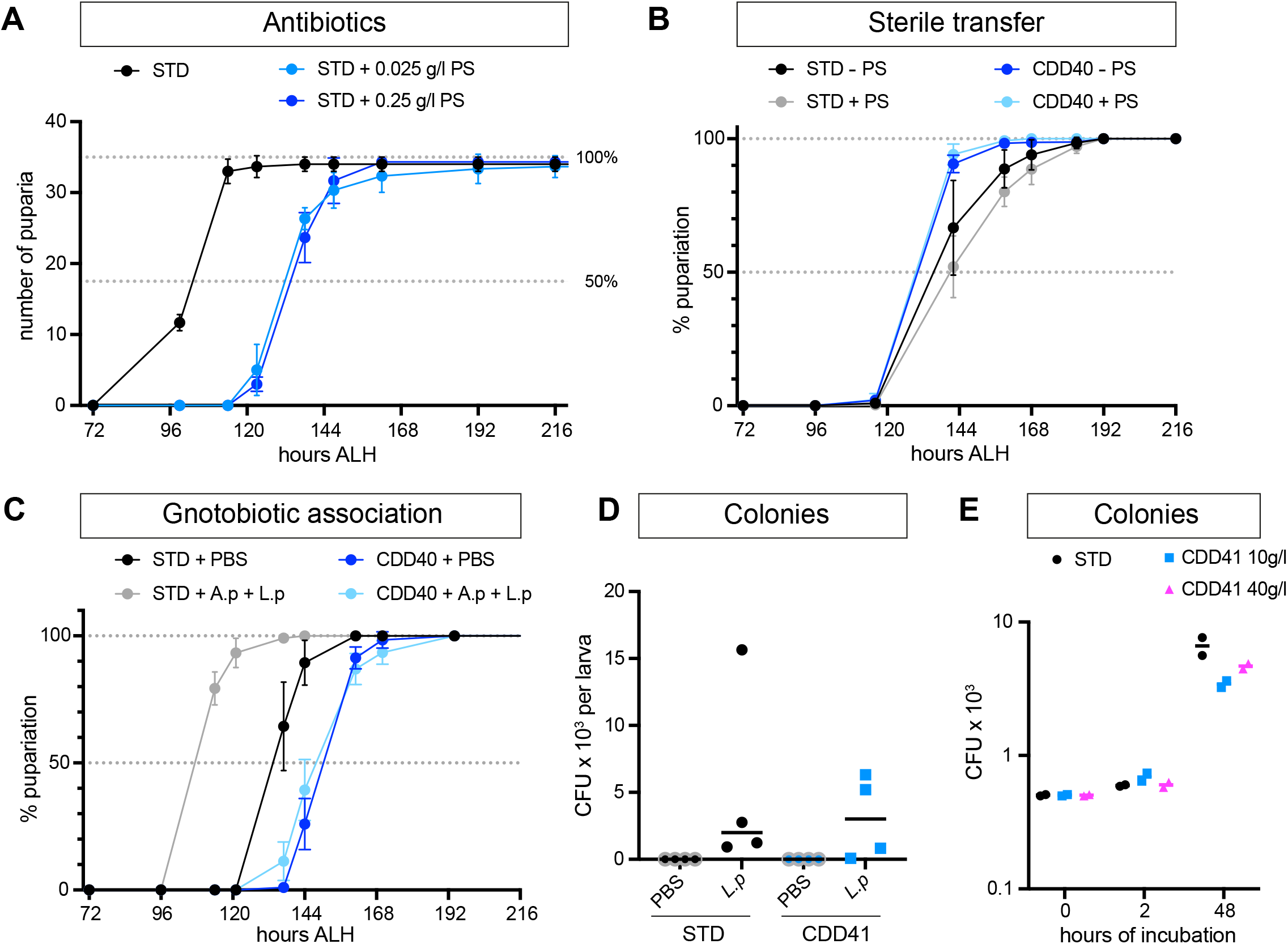
Microbiota accelerate larval growth on STD but not HoldFast. A) Time-to-Pupariation (TTP) of *w^1118^ iso 31* larvae on STD with or without Penicillin and Streptomycin (PS) at the indicated concentrations. Antibiotics delayed pupariation on STD by ∼30 hr (TTP_50_ is ∼ 132 hr). B) TTP of sterile-transferred *w^1118^ iso 31* larvae on STD or CDD40 with or without PS (0.25 g/l). Under sterile-conditions, TTP_50_ on CDD is not altered whereas, on STD, it is greatly increased to ∼134 hr compared to non-sterile transfer. PS does not substantially alter TTP_50_ on sterile CDD or STD, indicating it acts as an antibiotic not a general developmental inhibitor. C) TTP of wild-type *Oregon R* larvae sterile-transferred to STD or CDD40 without antibiotics and inoculated with PBS or *Acetobacter pomorum^WJL^* and *Lactiplantibacillus plantarum^NC8^* (A.p + L.p). Gnotobiotic association rescues TTP on STD but not CDD. D) Colony-forming units (CFUs) from individual *w^1118^ iso 31* larvae sterile transferred to STD or CDD41 (both without antibiotics), inoculated with or without *Lactiplantibacillus plantarum^NC8^* (L.p). Variable numbers of CFUs can be isolated from larvae exposed to L.p on both diets. E) CFUs (log_10_ scale) from sterile vials (no PS), with no larvae, of CDD41 containing 10 or 40 g/litre of sucrose or STD vials, inoculated with *Acetobacter pomorum^WJL^* and *Lactiplantibacillus plantarum^NC8^* (A.p + L.p). Values at 0 hr were determined prior to inoculation and so are identical between the three diets. CFUs were determined on MRS plates, which primarily support L.p not A.p.

The antibiotic and sterile transfer experiments suggest that TTP_50_ on holidic diets might be further improved if functional microbiota could be sustained. We therefore measured the effects of gnotobiotic association of sterile transferred embryos with known commensal species. Consistent with a previous study^68^, innoculation of sterile transferred *Oregon R* embryos on STD with *Acetobacter pomorum^WJL^* and *Lactiplantibacillus plantarum^NC8^* (hereafter *A.p* and *L.p*) shortened TTP_50_ to a value close to that obtained in the presence of the full *Drosophila* microbiome (**Figure 5C**, compare with **Figure 5A**). In contrast, gnotobiotic association of *Oregon R* on CDD40 with *A.p* and *L.p*, *A.p* alone, or A.p supplemented with one of its preferred nutrient sources (lactate), produced no significant improvement in TTP (**Figures 5C** and **S6B**). Similarly, gnotobiotic association of *w*^*1118*^ *iso 31* embryos with *A.p* and *L.p* did not shorten TTP_50_ on CDD40 and, in some experiments, even moderately delayed it (**Figure S6C**). This negative impact on the TTP_50_ of *w*^*1118*^ *iso 31* larvae was rescued by supplementation with inulin or lactulose but these prebiotics were unable to close the TTP_50_ gap between CDD40 and STD under conditions of gnotobiotic association (compare **Figures S6C** and **S6D**). Furthermore, the TTP_50_ of non-sterile embryos transferred to CDD was very similar with or without PenStrep, indicating that developmental speed on CDD is not significantly altered by the conventional microbiome present in our lab stock (**Figure S6E**). One possible reason why commensal bacteria may improve the speed of development on STD but not on CDD formulations is that they do not thrive on the holidic diet. However, colony forming units were recovered from sterile-transferred larvae associated with *L.p* on both STD and CDD41 and, even in the absence of larvae, *A.p* and/or *L.p* can multiply in two days ∼8-fold on CDD41, compared to ∼13-fold on STD (**Figures 5D** and **5E**). Hence, commensal members of the microbiota can thrive on both STD and CDD diets but they can only function to accelerate TTP on STD. Together, these results indicate that the pace of development on CDD formulations is not dependent upon microbiota. They also demonstrate that functional microbiota are likely to account for most, if not all, of the remaining differences in developmental pace between STD and CDD41.

### Optimal holidic diets for larval development and adult lifespan are different

The results thus far raise the important question of whether or not an optimal holidic diet for larvae is also optimal for adults. For larval-optimised CDD41, we observed two adverse adult outcomes on adult *w*^*1118*^ *iso 31* flies. First, newly eclosed flies tended to stick to the food surface and die (**Figure 6A**). And second, for male flies reared to eclosion on STD, lifespan was substantially shorter on CDD41 (32 days median lifespan) than on STD (60 days median lifespan) (**Figures S7A** and **S7B**). To circumvent the first of these limitations, a less sticky version of CDD41 was formulated in which the concentration of 2-hydroxyethyl agarose (low-melting point agarose) was raised by 50%, fully ameliorating the early adult mortality hazard (**Figure 6A**). In addition, we doubled the concentration of sucrose, which we showed before to be compatible with fast development and growth, to create CDD48. When adult survival was re-tested on CDD48, using *w*^*1118*^ *iso 31* males and females, median lifespan was measured at 45 days in both sexes (**Figure 6B**). This indicates that median lifespan on CDD48 is considerably longer than on CDD41, although still 10-15 days less than on STD. To test the effect of CDD48 on fecundity, we quantified egg deposition by adult females and found this to be substantially increased compared to STD, either with or without PenStrep (**Figure 6C**). We conclude that CDD48 performs well for adult fecundity, but other holidic diets specifically formulated for adults can support greater longevity. It is known that lifespan can be increased by lowering the dietary protein-to-carbohydrate ratio or by dietary restriction (food dilution) but both nutritional manipulations also tend to decrease fecundity^22,69,70^. Given this, it is perhaps not surprising that CDD48 is suboptimal for longevity as, in order to support rapid larval development, it has an amino acid content and an amino acid-to-carbohydrate ratio higher than yeast-based and most other holidic diets.

**Figure 6:**
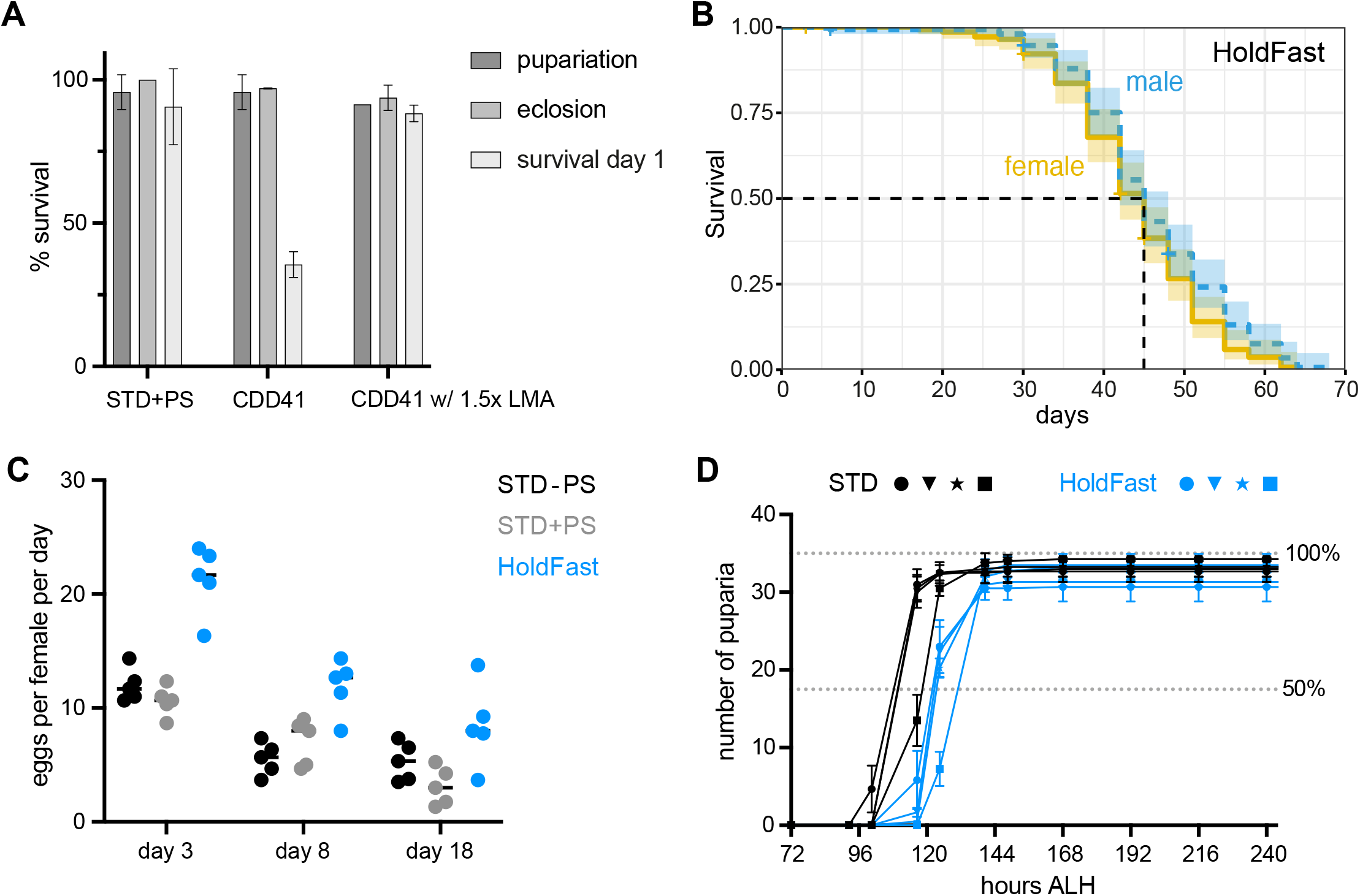
HoldFast (CDD48) supports a higher fecundity but a shorter lifespan than STD. A) Quantification of survival of *w^1118^ iso31* animals on diets with antibiotics (STD+PS or CDD41+PS) with 1% or 1.5% 2-hydroxyethyl agarose). Percentage survival was quantified at three stages: pupariation, adult eclosion and 24 h after eclosion. B) Kaplan-Meier survival curve of *white iso31* males and females on HoldFast (CDD48). Median lifespan is ∼45 days for both sexes. Shading indicates 95% confidence interval. C) Female fecundity on HoldFast (CDD48) or STD with or without antibiotics (± PS), measured at day 3, 8 and 18 after eclosion. HoldFast gives more eggs per female per day than either STD diet at all three timepoints. D) Time-to-Pupariation (TTP) of *w^1118^ iso 31* larvae on STD (-PS) or HoldFast (CDD48). Independent replicates (N=4 experiments) paired for STD and CDD48 (indicated with identical symbols).

Finally, we tested the ability of CDD48 to support larval development, finding that TTP_50_ values are very similar to those obtained for CDD41 and less than a day longer than for STD with no PenStrep (**Figure 6D**). Hence, the two improvements to CDD41 that yielded CDD48 are able to increase adult lifespan without a detectable larval tradeoff. We henceforth rename CDD48 as HoldFast (**<u>Hol</u>**idic **<u>d</u>**iet for **<u>Fast</u>** development). A direct comparison between holidic diets indicates that HoldFast (with or without antibiotics) represents a substantial improvement over the FlyAA diet^36 34^, both in terms of the survival and the TTP of *w^1^*^18^ *iso31* and *Ore R* larvae (**Figures S8A** and **S8B**). One contributing factor to the lower performance of FlyAA appears to be micronutrients as doubling their concentration substantially shortened TTP_50_ and, at least for *Ore R*, also improved survival (**Figures S8A** and **S8B**). In summary, for two different genetic strains, developmental speed and survival are substantially greater on HoldFast than FlyAA.

## Discussion

This study reports, for the first time to our knowledge, a holidic medium that supports a the rapid rate of development characteristic of *Drosophila*. HoldFast is composed entirely of high-purity chemicals and is capable of sustaining fast larval development, growth and survival at comparable levels to standard yeast-based diets. Multiple macro- and micronutrient improvements were combined during the larval optimisation of HoldFast. These include the use of amino acid and vitamin profiles based on autolysed yeast, an increase in vitamin concentrations, and the addition of the plant sterol sitosterol. To improve the performance of HoldFast for adult lifespan without a noticeable larval tradeoff, the concentrations of sucrose and 2-hydroxyethyl agarose were increased. We now discuss the co-optimisation of individual macro- and micronutrients and their effects on developmental speed, growth and survival. We then consider the role of the microbiome in holidic versus oligidic diets, and highlight some uses and limitations of HoldFast for *Drosophila* research.

### Amino acid ratios optimised for fast development

HoldFast was developed from a holidic diet with poor larval performance (CDD22) via multiple optimisation and co-optimisation steps designed to speed up development. This process was guided by our detailed nutritional analysis of a standard yeast-based diet and it built upon many previous *Drosophila* dietary studies. In this way, we successfully identified ratios of dietary amino acids that accelerate development towards the fast pace supported by standard yeast-based diets. Consistent with a previous study^36^, we found that dietary amino acid ratios that match the *in silico* translated *Drosophila* exome tended to perform better than strongly mismatched ones. We also showed that, for yeast, the amino acid profile of the *in silico* translated genome turns out to provide a rough proxy for the proteome, which may well also be the case for *Drosophila* and other species. Importantly, however, an amino acid profile matching the composition of a yeast-based diet performed just as well as one that matched the translated *Drosophila* exome. In line with these findings, unsupervised PCA showed that the amino acid ratios of translated exomes from many diverse species, including *Drosophila* and yeast, tend to cluster together with diets of high larval performance but away from those of poor larval performance. More generally, our analysis also suggests that exome matching a diet to the species it is designed for may not always produce a better performance than matching it to another species.

### Non-essential roles of dietary fatty acids and carbohydrates

Fatty acids are not an essential component of *Drosophila* holidic diets^33^ and most formulations lack them, including HoldFast. It is surprising that an optimal pace of larval development is possible when fatty acids are supplied entirely by the energetically costly process of *de novo* synthesis rather than via the diet. Genetic analysis showed that the fat body is a critical site for this *de novo* synthesis, mediated via fatty acid synthases (*FASN1* and *FASN2*). Hence, dietary supplementation with palmitic acid is sufficient to bypass the developmental requirements for fat body *FASN1*+*2* on HoldFast. These and other experiments support the conclusion that fatty acids are essential for development but they can be supplied either from the diet or via *de novo* synthesis in the fat body. The main components of HoldFast capable of contributing to fatty acid synthesis are amino acids and the sole carbohydrate ingredient - sucrose. We found that sucrose is not essential for larval survival on Holdfast but there is a “sweet spot” concentration that maximizes developmental speed. Supplementation experiments indicated that the addition of high levels of unsaturated free fatty acids to HoldFast strongly decreases survival. Nevertheless, some complex lipid mixtures such as lard or a balanced mix of saturated (stearic) and unsaturated (oleic) fatty acids are compatible with survival and fast rates of growth and development. This indicates the potential utility of HoldFast for studies of developmental metabolism under conditions with a dietary fatty acid input, akin to the case with yeast-based media.

### Co-optimisation and interdependence of macro- and micronutrients

We found that dietary amino acid ratios designed for developmental speed tended to come at the expense of developmental survival. However, survival on “fast” amino acid ratio diets could be fully restored by co-optimisation of the micronutrients in the CDD22 base diet. This striking finding strongly suggests that a fast pace of growth and development places additional metabolic demands on the micronutrient pool. More generally, it highlights that the developmental requirements for individual macro- and micronutrients are interdependent and so cannot be optimised in isolation. In particular, survival on a “fast” amino acid diet was greatly increased by raising the concentrations of choline and multiple B vitamins at the physiological bioactive ratios present in a yeast-based diet. Survival, as well as growth, was also improved by switching sterols from animal-derived cholesterol to plant-derived β-sitosterol and also by raising the overall sterol concentration. *Drosophila* converts dietary cholesterol into 20-hydroxyecdysone but dietary β-sitosterol is metabolised into another bioactive ecdysteroid, makisterone A^49,71^. We found that the inclusion of cholesterol with β-sitosterol did not compromise fast larval development on HoldFast and this sterol mix has the practical advantage of being compatible with existing ecdysteroid immunoassays, which specifically measure 20-hydroxyecdysone^72^.

### HoldFast is a new tool for studying developmental metabolism

HoldFast opens up new possibilities for studying the roles and requirements of individual macro- and micronutrients during *Drosophila* development. It can be combined with genetic analysis, as illustrated for *FASN1+2*, and also with metabolomics and stable isotope tracing. In addition, HoldFast is customisable and panels of variant diets could be used to apply nutritional geometry^8,9^ to micronutrient as well as macronutrient influences on developmental parameters. From a practical standpoint, the use of high purity chemicals in holidic diets such as HoldFast should provide more consistency from batch-to-batch than for yeast-based diets.

### Limitations

A limitation of HoldFast is that it is optimised for fast larval development not adult longevity. More generally, our findings and several previous studies argue that it is unlikely that a single “one size fits all” holidic diet can be simultaneously optimal for all larval and adult traits. Even though we were able to increase adult survival on our holidic diet without a detectable larval tradeoff, the median lifespan still remains shorter than with some yeast-based and holidic diets. Further experiments will be necessary to show whether or not dilution of HoldFast extends adult lifespan, akin to the known effects of dietary restriction with yeast-based diets. Another potential weakness of HoldFast is that rapid larval development on this medium appears to be independent of an input from commensal bacteria. This may limit usefulness for microbiota research but, in some contexts, it will be advantageous to use a less complex nutritional landscape without bacterial inputs that affect developmental speed. Other diets that are also high in proteins or free amino acids, such as an 8% yeast diet^63^, can support fast larval development in the absence of microbiota. This contrasts with less rich diets such as our own 2% yeast diet (STD) and several other oligidic diets, where *Acetobacter* and *Lactiplantibacilli* species such as *A. pomorum^WJL^* and *L. plantarum^NC8^* accelerate development^44,62,63,65,66,73,74^. On these diets, microbiota are thought to function at least in part by increasing the availability of nutrients such as amino acids and vitamins to the host^44,51,56,72^. Microbiota can also impact other properties of the diet such as moisture content^74^ and this could be relevant for larval development as well as for adult survival. Our results do, however, suggest that microbial effects, whatever their nature, are not rate limiting for larval growth and development on nutrient-rich HoldFast. Given that microbiota can nevertheless colonise HoldFast, it seems that their developmental benefit is far less obvious in this context than on our yeast-based diet. In future, it will be interesting to determine whether specific microbial contributions to larval development can be studied by diluting one or more components of HoldFast.

## METHODS

### Drosophila melanogaster strains and diets

A *Wolbachia*-free isogenized *Drosophila melanogaster* strain of *white*^*1118*^ *iso31* (Refs 13,75) or an *Oregon R* wildtype strain^76^ were used. *D. melanogaster* strains were maintained on a 12 hr light: 12 hr dark cycle at 25°C and 60 % relative humidity. For stock maintenance and for yeast-based diet larval experiments, Standard Diet (STD) was used, which contains (w/v): 6.63 % cornmeal, 5.85 % Glucose, 2.34 % autolysed yeast, 0.195 % Nipagin (methyl paraben), 0.00078 % Bavistin (carbendazim), 1.95 % Ethanol (v/v), 0.7 % Agar^13^. Holidic diets also contain the same concentrations of Nipagin and Bavistin. Holidic diets also contain penicillin and streptomycin (PenStrep, PS) unless otherwise indicated. STD does not contain either antibiotic unless otherwise indicated. In those cases where Penicillin and Streptomycin were added to STD or to CDD this was at 0.25g of each per litre. For fat body expression we used *apoLpp-Gal4*^77^ and for gut enterocytes *NP1-Gal4* (also called Myo1A-Gal4, P{GawB}Myo31DF^NP0001^, Ref 78). A double UAS-RNAi stock targeting both *FASN1* and *FASN2* (TRiP.HM05141 and P{KK106426}VIE-260B) was used for FASN knockdown.

### Holidic diet preparation

Holidic diet formulations are listed in **Supplemental Table 1**. All chemically defined diets including HoldFast were prepared in essentially the same way and contained the same total mass of amino acids, 25g per litre. This amount is identical to that used in two previous holidic diets^33,37^, and ∼15% higher than that used in the Piper *et al.* 2013 holidic diet^34^. All ingredients of the “Amino acid mixture” (**Supplemental Table 1**) were weighed out and added to a glass beaker containing ultrapure water (Milli-Q Water Purification System, Merck Millipore). Tyrosine was separately pre-dissolved in 2.8 M KOH. Then, the “Sterol mixture” was prepared by dissolving sterols in a solution of Nipagin (methyl 4-hydroxybenzoate) and Bavistin (carbendazim) dissolved in ethanol (100g Nipagin and 0.4g Bavistin in 1 litre of pure Ethanol), added to the amino acid mixture and the pH of the whole mix adjusted to 6.5 with NaOH. The solution was then poured into an automatic cooker (Mediaclave 10, Integra Biosciences) and the agarose slowly added while stirring. The cooking programme heated to 100°C, held this temperature for 3 minutes and then cooled down to 50°C. This mix was then combined with the vitamin mix, pre-warmed to 50°C after being prepared separately with all components added to a beaker with ultrapure water and adjusted to pH 6.5 with NaOH/HCl. Thoroughly mixed diet was then dispensed manually via a plastic pipette into plastic vials.

For lipid supplementation experiments, CDD was melted in a water bath at 70°C, cooled to ∼50°C, and mixed thoroughly with liquid fats (corn oil, oleic acid, linoleic acid, palmitoleic acid) or solid fats (lard, coconut oil, palmitic acid, stearic acid). After cooling back into a gel, small fat droplets or precipitates were visible, presumably because CDD does not contain an emulsifier. The lipid concentrations used (0.5, 5 or 50 g/l) were centred on 5g/l, the amount estimated in STD from the 4-6% fatty acid content of cornmeal listed in various nutritional databases.

### Analysis of larval developmental parameters

Unless otherwise stated, first instar (L1) larvae of both sexes were transferred from grape juice-agar plates to vials containing different diets at 0-2 h after larval hatching. Transfer was carried out with a single blunted, bent forceps arm, acting like a tiny shovel. For TTP graphs, numbers of puparia were counted 1-3 times daily. Survival to the puparial stage was calculated as a percentage from the total number of puparia formed divided by total number of L1 larvae transferred. The number of larvae transferred per vial, typically 35, is marked on TTP graphs as 100%. Adult survival was calculated as a percentage from the number of empty pupal cases divided by the total number of puparia formed. Adult flies were separated into males and females and weighed individually on an MPE6.3P ultra-microbalance (Sartorius) at 2-4 days after transfer of eclosed individuals from holidic diet to STD vials. For germ-free experiments, embryos were collected from grape juice-agar plates in a small sieve, rinsed thoroughly with ultrapure water and then surface-sterilised with bleach (20% sodium hypochlorite solution). Bleaching was monitored under a dissecting scope and embryos were constantly agitated to ensure complete sterilisation. Embryos were then extensively rinsed with ultrapure water to remove bleach and transferred to vials using a brush pre-sterilised with ethanol and UV light exposure. Using this protocol, no microbial colonies were recovered from larvae or from vials.

### Lifespan and fecundity analysis

Fly lifespan was assessed using a standard “once mated” methodology^13^. Briefly, embryos were collected from large grape juice-agar plates, washed and distributed with a pipette to bottles with STD diet to achieve low-density conditions for larval growth. Upon eclosion, adults were collected and groups of 10 males and females mated for 1-3 days in STD vials. Vials were then pooled, separated by sex and distributed to each diet in groups of 15 per vial. A total of 150 flies (10 vials) were used per condition. Dead flies were scored every time flies were passed to a fresh vial, which occurred on average every three days.

For fecundity analysis, freshly eclosed virgin females were mated with males for three days on STD as for lifespan analysis, then groups of 4 females were housed on the respective diets. For the measurement, females were passed to a fresh vial and allowed to lay eggs for around 18 hours. Flies were then passed on again and all eggs laid were counted and reported as eggs per female and day (24 h). Five vials of four females were used per diet.

### Microbiota manipulation and analysis

Whole larval extracts were prepared by lysing an individual larva in 100µl PBS in a Precellys Evolution homogeniser using the “soft” setting. The extract was diluted 1:10 in PBS and 50µl plated on De Man–Rogosa–Sharpe (MRS) agar with 2g/L bacto-agar or mannitol agar (Bacto peptone 3 g/L, yeast extract 5 g/L, D-mannitol 25 g/L, bacto-agar 2 g/L) according to previous studies^44,68^. Agar plates were incubated at 30°C for 30 hr and colonies manually counted. Gnotobiotic association was performed essentially as described^44,68^. The *Acetobacter pomorum^WJL^* strain^62,79^ was grown as liquid culture in Mannitol broth (Bacto peptone 3 g/L, yeast extract 5 g/L, D-mannitol 25 g/L) at 30°C and shaking (180rpm). The *Lactiplantibacillus plantarum^NC8^* strain^62,79^ was grown as liquid culture in MRS broth at 30°C without shaking. Liquid cultures were collected by centrifugation, washed with PBS and then diluted appropriately in PBS before addition to vials with embryos. *Drosophila* embryos were collected on plates, washed thoroughly and sterilised with sodium hypochlorite before being distributed to sterile CDD or STD vials using a sterile brush. Sterile embryos were associated with cultured bacterial strains using ∼10^7^ colony forming units (CFUs) of either *Acetobacter pomorum^WJL^* or *Lactiplantibacillus plantarum^NC8^*, or a mix of ∼10^7^ CFUs of each bacterial species per vial of 10ml diet.

### LipidTOX staining

Fat bodies from stage-matched larvae were dissected and fixed for 25 min with 4 % paraformaldehyde in phosphate buffered saline (PBS). After washes with PBS, samples were incubated overnight with 1:500 LipidTOX Deep Red to stain neutral lipids and 1:2000 Hoechst 33342 DNA stain. After further PBS washes, fat bodies were mounted in Vectashield (Vector Laboratories) on microscope slides with a hole-punched tape spacer and imaged on a Leica Stellaris-8 inverted confocal microscope using the same acquisition settings for each sample.

### Analysis of translated exomes

The amino acid compositions of the translated exomes of 18 selected species were derived from their proteomes via <u>Proteome-pI</u>^80^. This approach is based on annotated proteins in the <u>UniProt</u> database and yields similar results to those obtained via *in silico* translation of exome sequences^36^. Species were selected to span all domains of life and to include the standard model organisms. For PCA analysis, data was centred and autoscaled before analysis using the R package stats (version 4.2.2) and plotted using the R package <u>factoextra</u> (version 1.0.7). All analysis was performed using <u>R</u> version 4.2.2. The Top25 expressed proteins of yeast (*S. cerevisiae*) were manually curated from the <u>Saccharomyces Genome Database</u>, based on microscopy or flow cytometry measurements: Eno2p, Ssa2p, Tdh3p, Pdc1p, Tef1p, Fba1p, Pgk1p, Tef2p, Hsp26p, Tdh2p, Tpi1p, Rpl36Ap, Rpl36Bp, Ald4p, Pdc1p, Ssa1p, Ahp1p, Eno1p, Asn2p, Yef3p, Rps26Bp, Rps26Ap, Rnr1p, Rnr4p, Hsc82p.

### Measurements of dietary amino acids and bioactive vitamins

Individual amino acids in STD diet as well as its main components (autolysed yeast, cornmeal, agar) were quantified in duplicate samples by AltaBiosciences (Birmingham, UK). Amino acids were analysed by acid hydrolysis using 6N hydrochloric acid for 24 h at 110°C, followed by subsequent lyophilization, resuspension and separation via ion exchange chromatography and detection with ninhydrin. Acid hydrolysis provides values for Glu+Gln and for Asp+Asn but not for the corresponding individual amino acids. Cysteine was separately quantified following oxidation by performic acid followed by the same procedure as for the other amino acids. Tryptophan was quantified using alkaline hydrolysis (2.5 g barium hydroxide in 2ml H_2_O for 16 h at 110°C), followed by precipitation with sodium sulphate and separation through ion exchange chromatography and detection with ninhydrin. Bioactive vitamin levels were determined by the <u>Institut für</u> <u>Produktqualität</u> (ifp, Berlin, Germany) using <u>VitaFast</u> microbiological bioassays for all B vitamins and by <u>LC-MS/MS</u> for total choline. VitaFast assays quantify bioactive forms of vitamins via the growth of bacteria of the *Lactiplantibacillus* genus, auxotrophic for the vitamin being measured, and are calibrated with pure standards. Single samples of autolysed yeast and cornmeal and duplicates of cooked STD diet were analysed.

### Hemolymph extraction and NMR metabolomics

Hemolymph was extracted and analysed by NMR using the paired volume determination with two standards (pVDTS) method, essentially as described^81^ but with the following modifications. Groups of 10 to 12 third-instar larvae were immersed in a drop of ice-cold 0.9% w/v saline solution containing 20mM not 25mM sodium formate-13C and the 7.5µl blank from unopened larvae was removed after ∼15 seconds not 1 minute of immersion. Paired blanks (from unopened larvae) and hemolymph samples (from opened larvae) were filtered through a 0.22µm spin filter (Millipore) with 200µl deionized water containing 50µM sodium-4,4-dimethyl-4-silapentane-1-sulfonate (DSS). 200µl of flow-through was transferred to glass vials containing 500µl methanol and 250µl chloroform. Phase separation was achieved via the addition of 250µl chloroform, then vortexing and the addition of 250µl deionized water, further vortexing and brief microcentrifugation. The upper, polar phase was collected, dried under vacuum, resuspended in 170µl PBS in deuterium oxide (D_2_O) and 160µl loaded into Bruker Biospin 3mm NMR glass tubes. Spectra were acquired on a Bruker Avance III 700Mhz NMR spectrometer, using <u>Bruker</u> <u>TopSpin software</u> v3.6 and the standard pulse sequence noesypr1d. Spectra were analysed and quantified in <u>Chenomx NMR Suite</u> v9 using the build-in library and a custom spectrum for 13C-formate. Hemolymph concentrations of amino acids were calculated using the pVDTS formula as described^81^.

## QUANTIFICATION AND STATISTICAL ANALYSIS

Statistical analysis was applied to dietary formulations that showed improvements (see Design). These were prepared and tested independently via multiple experiments conducted on different days using larval collections from separate parental cages. The initial holidic diet (CDD22) and all major improvements (CDD33, 40, 41, 48) were prepared independently three or more times. For other formulations, three to five replicate vials were tested using larval collections usually from the same parental fly cage. TTP traces in graphs show average numbers of puparia between replicate vials and error bars show +/- one standard deviation. Statistical analysis of adult weights was performed in GraphPad <u>Prism</u> (version 10) or R using one-way ANOVA with Welch’s correction and Dunnett’s T3 test for multiple comparisons to the control/initial condition or with pairwise comparisons. For lifespans, data were analysed using the R <u>‘survival’</u> <u>package</u> (version 3.5-5) and plotted using the R <u>‘survminer’ package</u> (version 0.4.9). Graphs depict Kaplan-Meier survival probability, shading indicates the 95% confidence interval, and p values were determined using the log rank test.

## Acknowledgements

We thank François Leulier for bacterial strains and the late Suzanne Eaton for the *apoLpp-Gal4* stock. Fly stocks were also obtained from the <u>Bloomington *Drosophila*</u> <u>Stock Center</u> (BDSC, NIH P40OD018537), the <u>Vienna Drosophila Research Centre</u> (VDRC) and the <u>Kyoto Stock Centre</u>. We acknowledge technical assistance from the Crick Media Preparation Team and Paul Driscoll in the Crick Metabolomics STP for acquisition of NMR spectra. We also thank Andrew Bailey, Elisabeth Kamper and Clare Newell for advice and comments on the manuscript. Figure 1A was created with BioRender.com. This work was supported by the Francis Crick Institute, which receives its core funding from Cancer Research UK (CC2101), the UK Medical Research Council (CC2101) and the Wellcome Trust (CC2101). It was also supported by an Wellcome Investigator Award to APG (104566) and an EMBO ALTF to V.G (543-2022). For the purpose of Open Access, the authors have applied a CC BY public copyright licence to any Author Accepted Manuscript version arising from this submission.

## Author contributions

Conceptualization, S.S. and A.P.G.; Methodology, S.S. and A.P.G.; Investigation and formal analysis, S.S., V.G., L.L., V.T. and A.W.; Writing – Original Draft, S.S and A.P.G; Writing – Review & Editing, S.S, V.G., L.L., V.T. and A.P.G.; Funding Acquisition, V.G. and A.P.G; Supervision, T.H. and A.P.G.

## Supplemental information

**Supplemental Document S1**

**Figure S1:**
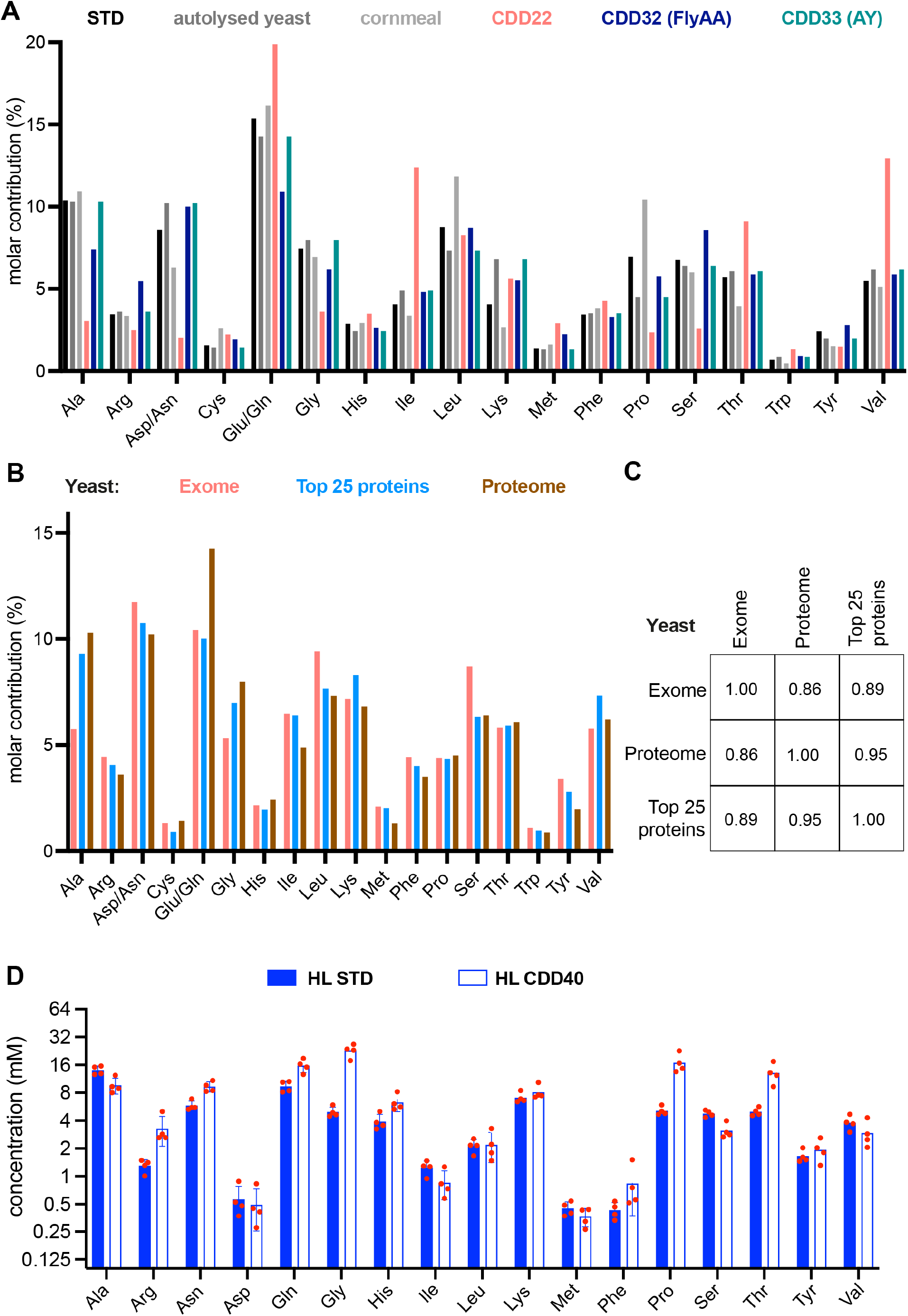
Amino-acid profiles of diets, dietary components, yeast and hemolymph. A) Relative (%) molar contributions of individual amino acids in STD diet, autolysed yeast (AY), cornmeal and three CDD22-based diets. The amino acid profiles of CDD32 (FlyAA) and CDD33 (AY) are based on the *in silico* translated *Drosophila* exome (Piper et al. 2017) and the composition of autolysed yeast respectively. In STD, AY and cornmeal, chemical hydrolysis does not distinguish between Glu and Gln, or between Asp and Asn, so values are plotted for the sum of both (Glu/Gln or Asp/Asn). B) Relative (%) molar contributions of individual amino acids to the yeast (*Saccharomyces cerevisiae*) exome, the top 25 most highly expressed proteins, and the proteome (as determined by chemical hydrolysis of AY in this study). As in A, values are plotted for the sum of Glu/Gln and for Asp/Asn. C) Spearmann’s rank correlation coefficients of the amino acid profiles of the yeast (*Saccharomyces cerevisiae*) exome, proteome and top 25 most highly expressed proteins. D) Hemolymph concentrations (mM) of amino acids from third-instar larvae raised on STD or CDD40. Note that several amino acids (especially Gly, Pro and Thr) are at higher hemolymph concentrations in larvae raised on CDD40 than on STD.

**Figure S2:**
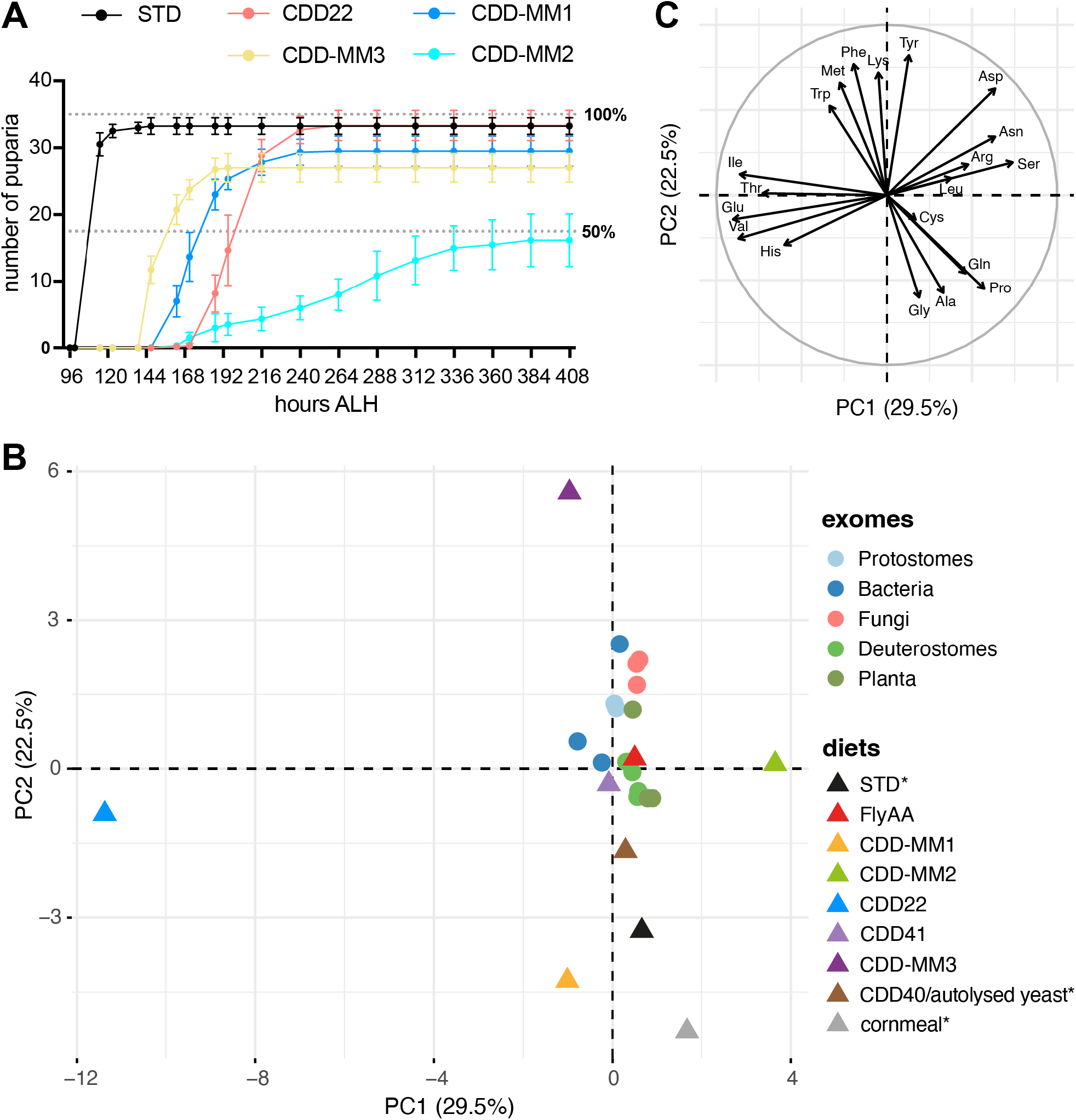
Performance and PCA analysis of STD and holidic diets. A) Time to pupariation (TTP) curves for STD, CDD22 and three amino acid-mismatched variant diets: CDD-MM1, CDD-MM2 (both from Piper et al.^36^) and CDD-MM3. B) Unsupervised Principle Component Analysis (PCA) of the amino acid profiles of *Drosophila* diets (diets) and annotated translated exomes derived from UniProt across the kingdoms of life (exomes). PC1 and PC2 respectively account for 29.5% and 22.5% of the variation. Individual species representing each domain of life are: Bacteria – *Escherichia coli, Lactobacillus plantarum, Helicobacter pylori*; Fungi – *Saccharomyces cerevisiae, Schizosaccharomyces pombe, Candida albicans*; Protostomes – *Drosophila melanogaster, Tribolium castaneum, Caenorhabditis elegans*; Deuterostomes – *Bos taurus, Gallus gallus, Homo sapiens, Danio rerio, Mus musculus, Oryzias latipes*; Planta – *Arabidopsis thaliana, Oryza sativa, Zea mays*. For STD, autolysed yeast and cornmeal, asterisks indicate that the values for Glu and Gln, as well as for Asp and Asn, were artificially split 50:50 between the acidic and amidic forms, as acid hydrolysis cannot discriminate between them. C) Loadings plot showing the influence of each amino acid in PC1 and PC2 space.

**Figure S3:**
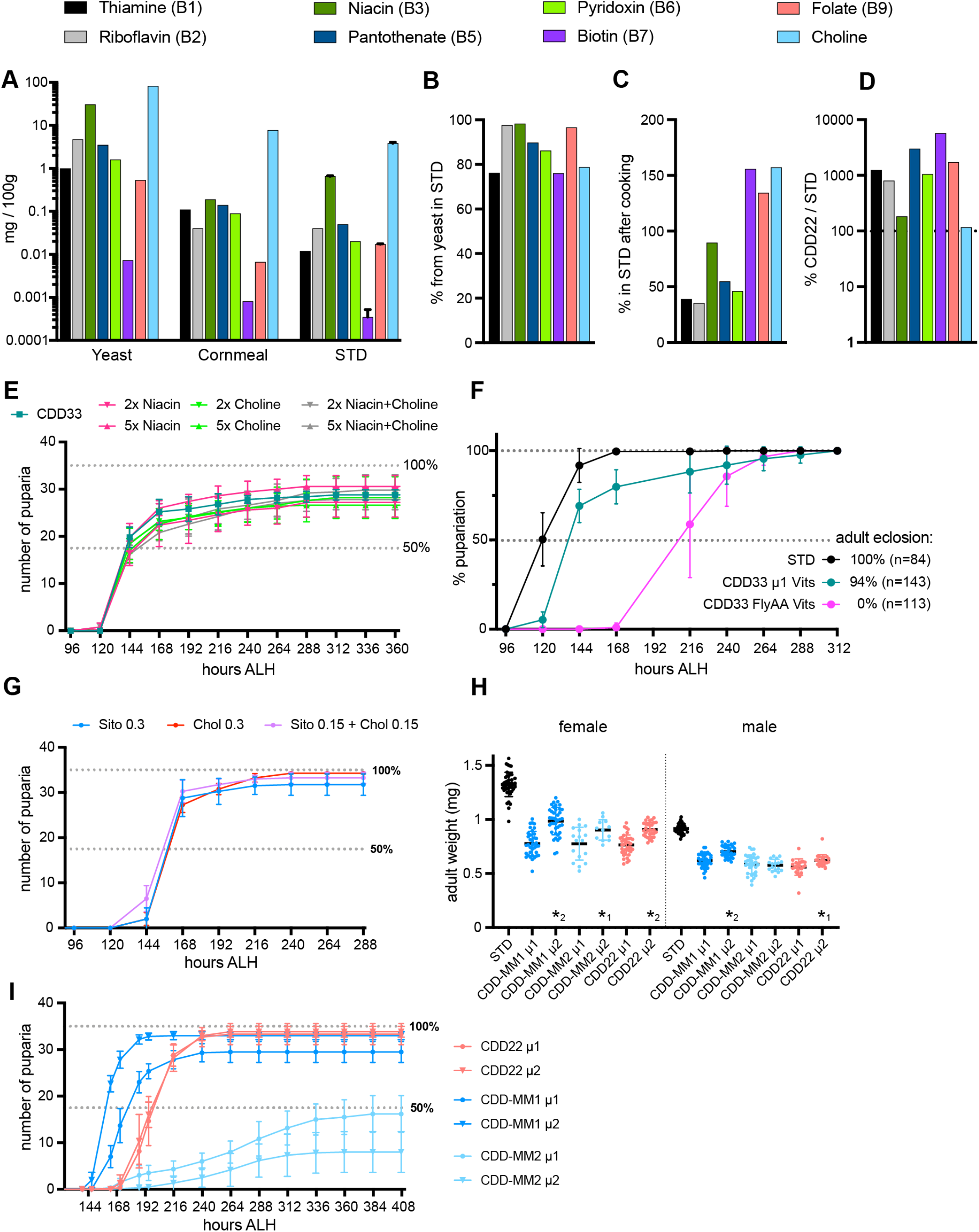
Vitamin and sterol effects on holidic diet performances. A) Bioactive vitamin concentrations (log2 scale, mg per 100g diet) measured in STD, yeast and cornmeal. Numbers of independent biological replicates are STD (N=2), yeast (N=1) and cornmeal (N=1). B) Calculated percentage of each vitamin in STD contributed by autolysed yeast. C) Percentage of each active vitamin remaining in STD after cooking, compared to the theoretical content before cooking. See text for details. D) Percentage of each vitamin in CDD22 compared to STD. All vitamins are higher, to varying degrees, in CDD than in STD E) Time-to-Pupariation (TTP) of control (*w^1118^ iso 31*) larvae on regular CDD33 containing 1x niacin and 1x choline, or with the increased amounts indicated. 5x Niacin showed a small increase in survival compared to CDD33 at 300 hr ALH. F) TTP of sterile-transferred control larvae on STD, CDD33 or CDD33-vits-FlyAA, formulated with the FlyAA vitamin mix. In the context of CDD33, the FlyAA vitamin mix greatly delays pupariation and is not compatible with adult survival. G) TTP of control larvae on CDD33 with either 0.3g/l Sitosterol, 0.3g/l Cholesterol, or 0.15g/l Sitosterol plus 0.15g/l Cholesterol. H) Adult male and female weights, at 2 to 4 days after eclosion, from the growth experiment in panel H. Physiologically optimised (µ2) micronutrients show a trend for increased growth with all three amino acid formulations, except for MM2 males. This contrast with the MM1-specific improvements of µ2 on developmental speed and survival (panel I). Statistical significance is indicated for pairwise comparisons between the µ1 and µ2 (a *1, p<0.05, A *2, p<0.0001). I) TTP of control larvae on CDD22 or the amino-acid mismatched diets MM1 and MM2, formulated with the original (µ1) or the physiological optimised (µ2) micronutrients (sterol and vitamins) mixes. Note that the µ1 data are the same as in Fig S1D. Optimised micronutrients (µ2) improved survival and developmental speed in the context of the mismatched amino-acid profile in MM1 but not those in MM2 or CDD22.

**Figure S4:**
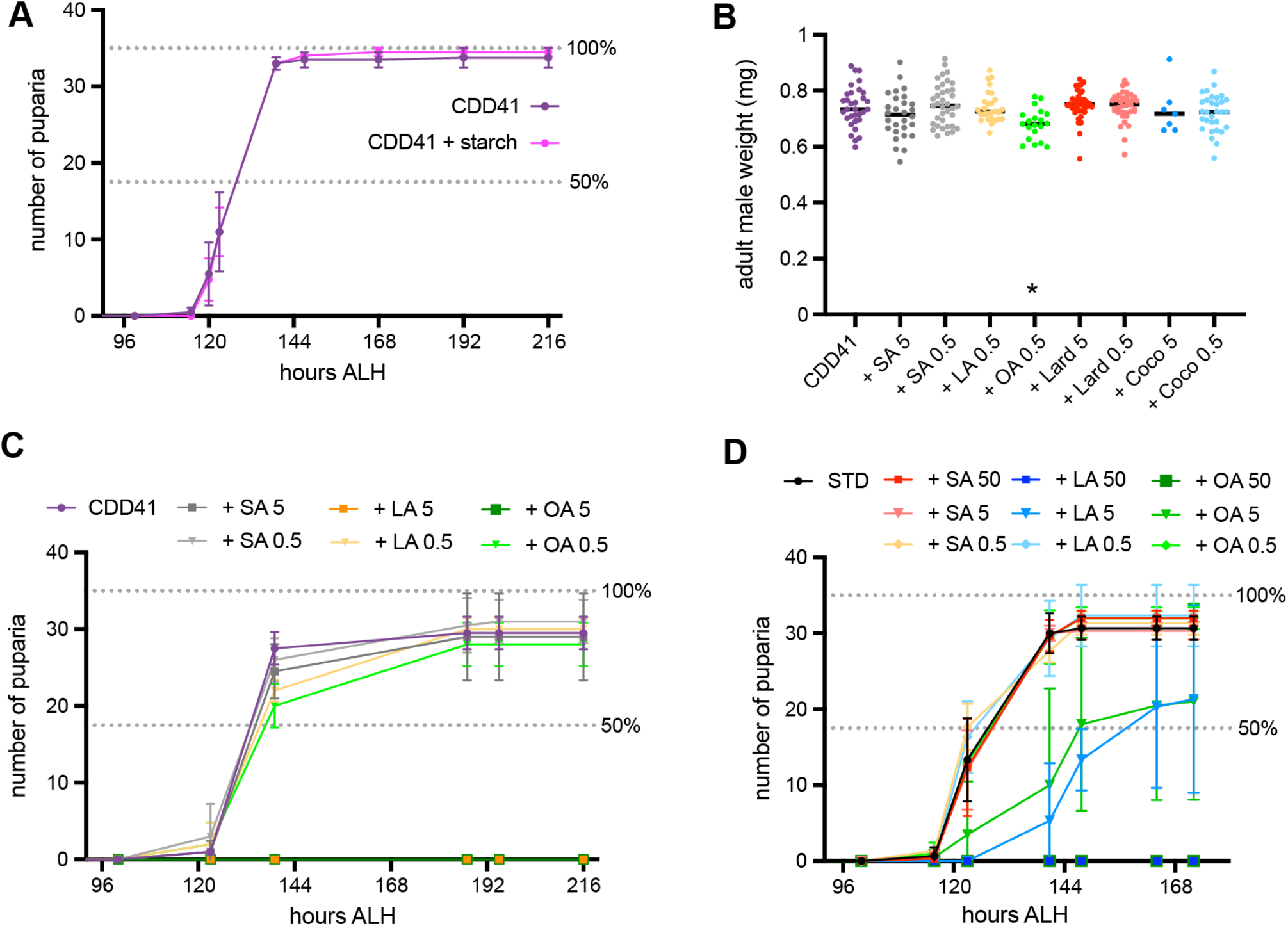
Developmental effects of starch or lipids in CDD and STD. A) Time-to-Pupariation (TTP) of *w^1118^ iso31* control larvae on CDD41. The addition of 40 g/l starch to CDD41 had no effect on developmental speed or survival. B) Adult male weights from the experiments in Figures 3C and S4C. Addition of stearic acid (SA), oleic acid (OA) or linoleic acid, or the natural fats coconut oil (Coco) or lard at 0.5 or 5 g/l did not substantially alter weights. There was a statistically significant weight decrease when 0.5 g/l OA was added to the CDD41 base diet (*, p=0.0096). C) TTP of *w^1118^ iso31* control larvae on CDD41 with the addition of stearic acid (SA), oleic acid (OA) or linoleic acid (LA) at 0.5 or 5 g/l. LA or OA at 5 g/l was larval lethal. D) TTP of *w^1118^ iso31* control larvae on standard diet (STD) with addition of SA, LA or OA at 0.5, 5 or 50 g/l. Linoleic or oleic acids are lethal at 50 g/l and they slow development and decrease survival at 5 g/l but are compatible with normal development at 0.5 g/l.

**Figure S5:**
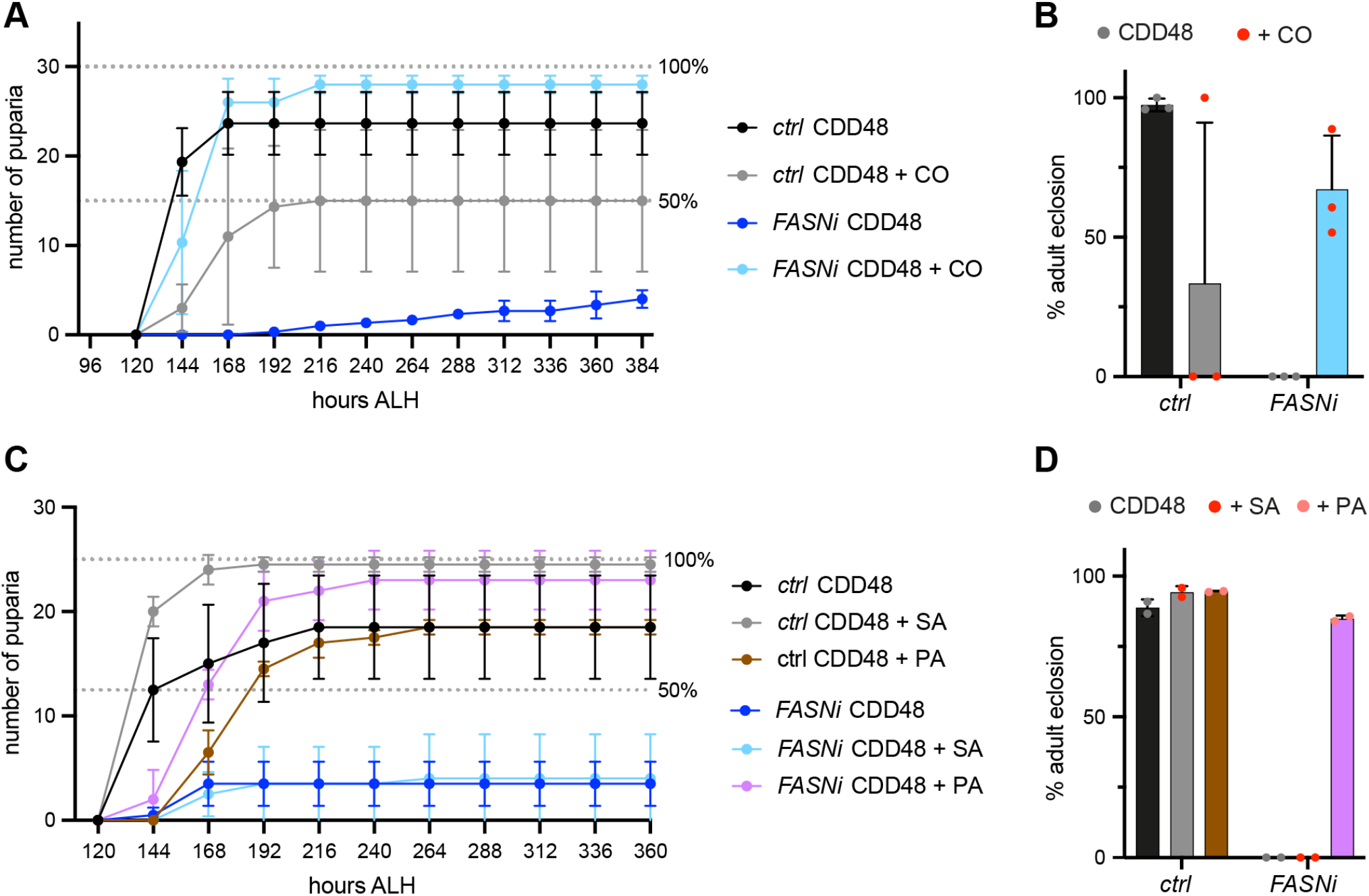
Palmitate or corn oil rescues the developmental lethality of fat body *FASN* knockdown. (A-B) Time-to-Pupariation (TTP) curves of larvae (A) and adult survival (% adult eclosion) (B) on CDD48 or CDD48 supplemented with 5 g/l corn oil (+corn). Larvae are control (*ctrl*) genotype (*Myo1A-Gal4* and *apoLpp-Gal4*), or fat body-specific FASN1+2 knockdown (*FB>FASNi* using *apoLpp-GAL4*). Corn oil was detrimental to *ctrl* larvae but rescued larval and adult viability of *FB>FASNi* animals. (C-D) TTP curves of larvae (C) and adult survival (% adult eclosion) (D) on CDD48 or CDD48 supplemented with 5 g/l of either stearic acid (+SA) or palmitic acid (+PA). Larvae are control (*ctrl*) genotype (*Myo1A-Gal4* and *apoLpp-Gal4*), or fat body-specific FASN1+2 knockdown (*FB>FASNi* using *apoLpp-GAL4*). Palmitic but not stearic acid rescued larval and adult viability of *FB>FASNi* animals.

**Figure S6.**
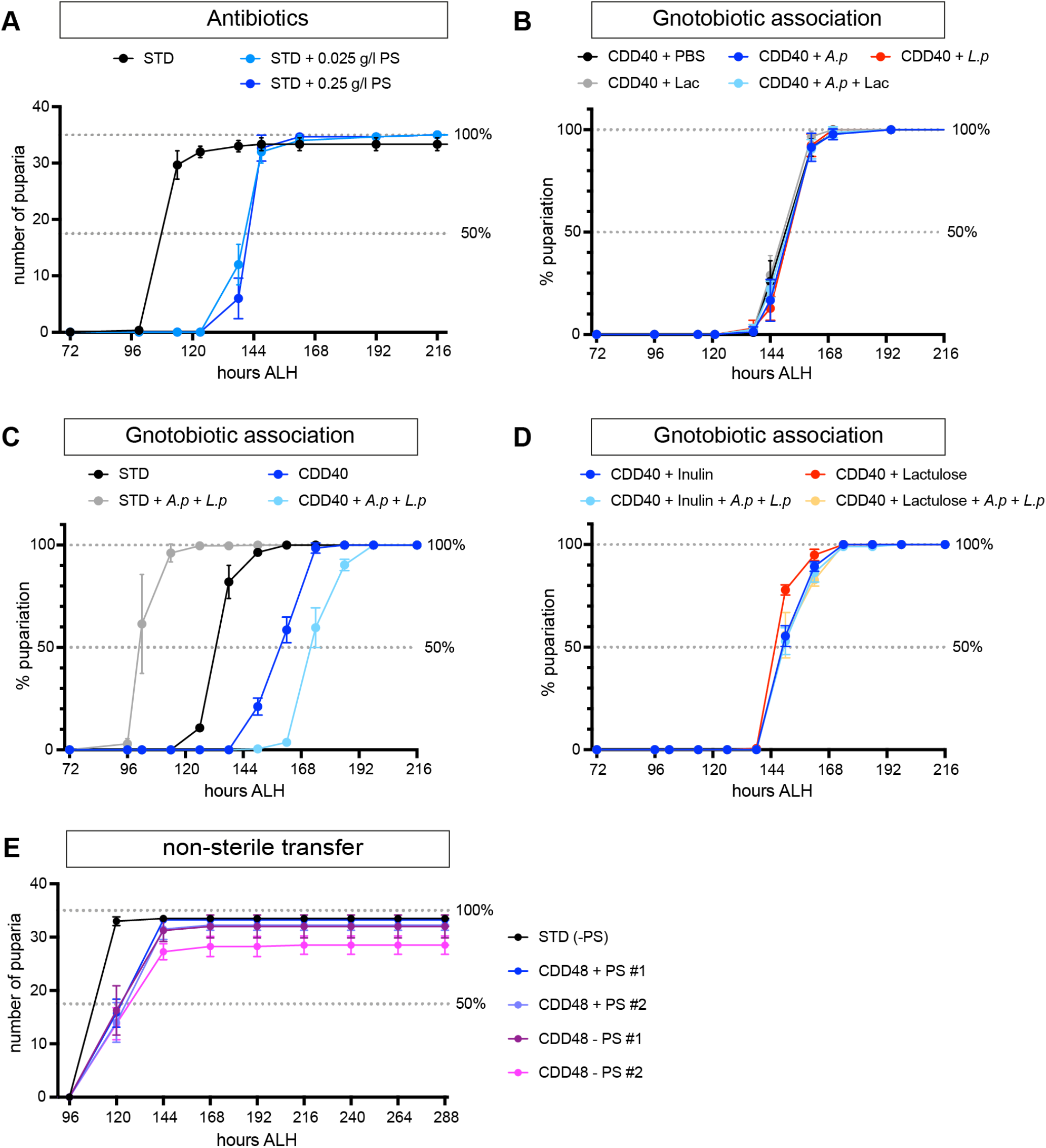
Gnotobiotic association accelerates larval growth on STD but not HoldFast. A) Time-to-Pupariation (TTP) of wild-type *Oregon R* (*OreR*) larvae on STD with or without Penicillin and Streptomycin (PS) at 0.25 or 0.025 g/litre. PS delays development on STD. The TTP50 of *OreR* larvae is ∼ 8 hr longer than for *w^1118^ iso31* larvae, both with or without antibiotics (compare with Fig 4A). B) TTP of sterile-transferred *OreR* larvae on STD or CDD40 lacking PS, with or without inoculation with *Acetobacter pomorum^WJL^* (*A.p*), *Lactiplantibacillus plantarum^NC8^* (*L.p*), or PBS. Supplementation with lactate (Lac), a nutrient substrate for *A.p*, did not alter TTP50. C) TTP of sterile-transferred *w^1118^ iso31* larvae on STD or CDD40 lacking PS but inoculated with *A.p* and *L.p,* or PBS. Commensal bacteria shorten TTP on STD but delay it on HoldFast. Fig S6C and S6D are from the same experiment but separated for clarity. D) TTP of sterile-transferred *w^1118^ iso31* larvae on CDD40 lacking PS but supplemented with prebiotics (inulin or lactulose), with or without *A.p* and *L.p* inoculation, or PBS. Inulin and lactulose had no substantial effect on TTP50 but comparisons with Fig S4D indicate that either prebiotic can rescue the developmental delay associated with *A.p* and *L.p.* Fig S6C and S6D are from the same experiment but separated for clarity. E) TTP of conventional (non-sterile) *w^1118^ iso31* larvae reared on STD lacking antibiotics (STD-PS) or CDD48 without or with antibiotics (CDD48-PS, CDD48+PS). Curves from two independent experiments (#1 and #2) are shown for CDD48. Antibiotics do not substantially alter TTP50 on CDD48.

**Figure S7:**
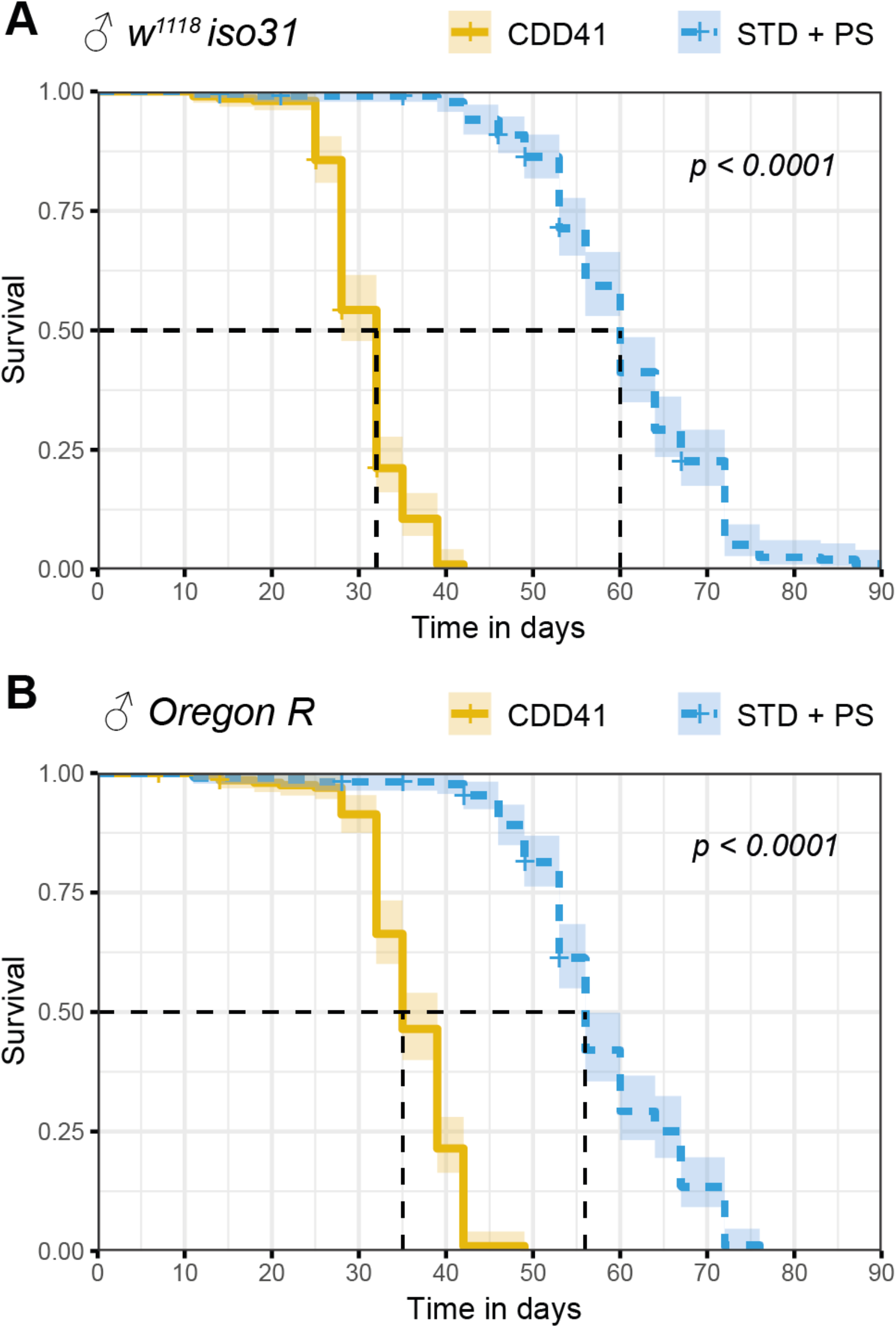
Lifespan on CDD41 is strongly reduced compared to STD. A) Kaplan-Meier survival curve of *white iso31* males on CDD41 or STD, both with antibiotics. STD males have a median lifespan of 60 days, while CDD41 supports a reduced lifespan of 32 days. B) Kaplan-Meier survival curve of *Oregon R* males on CDD41 or STD, both with antibiotics. STD males have a median lifespan of 56 days, while CDD41 supports a reduced lifespan of 35 days. Shading indicates 95% confidence interval and p values were determined using the log rank test

**Figure S8:**
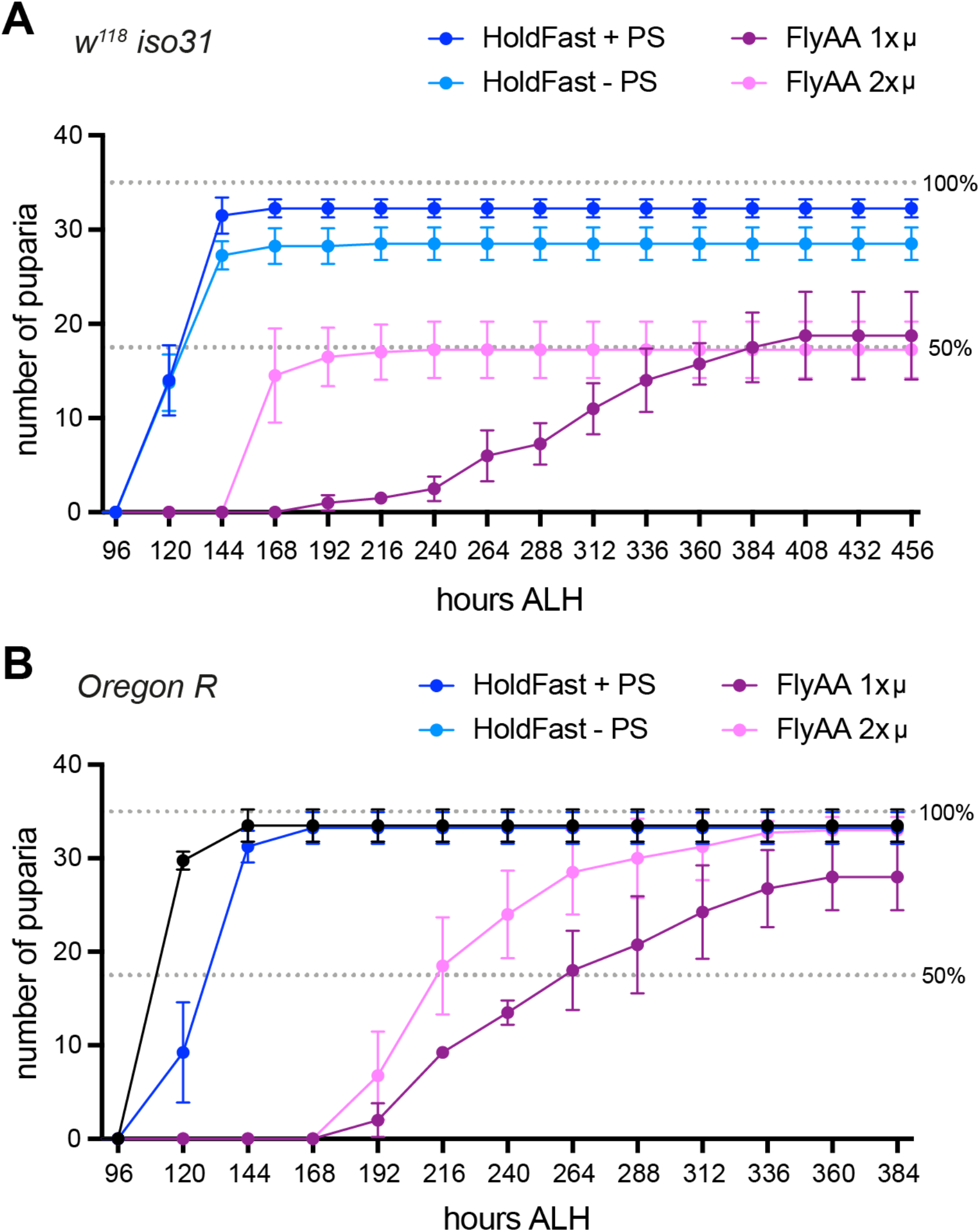
Comparison of HoldFast with FlyAA. A) Time-to-Pupariation (TTP) of conventional (non-sterile) *w^1118^ iso31* larvae reared on HoldFast (CDD48) without or with PenStrep (HoldFast-PS, HoldFast+PS) or reared on FlyAA diet (Piper *et al.* 2017) with original (1x) or double (2x) the concentration of micronutrients. HoldFast has double the survival and a TTP50 that is 5-6 days shorter than FlyAA. A two-fold increase in FlyAA micronutrients improved TTP50 without increasing larval survival to pupariation. HoldFast curves are from the same experiment as in Figure S6E (#2). B) TTP of conventional (non-sterile) wild-type *Oregon R* (*OreR*) larvae reared on HoldFast without or with PenStrep (HoldFast-PS, HoldFast+PS) or FlyAA diet with normal (FlyAA 1x) or double (FlyAA 2x) the concentration of micronutrients.

**Table S1: CDD/HoldFast formulations**

Excel file listing the amounts (in g) of all chemical components, the suppliers and the catalogue numbers for CDD22, CDD32(FlyAA), CDD33, CDD-MM1, CDD-MM2, CDD-MM3, CDD40, CDD41 and CDD48/HoldFast.

**Table S2: Dietary amino acid analyses**

Excel file listing the amino acid contents (g per 100g), determined by acid hydrolysis, from duplicate samples (#1 and #2) of yeast-based standard diet (STD), autolysed yeast, cornmeal, and agar.

