## Supplemental Table S1 for "A *Drosophila* holidic diet optimised for growth and development"

| Ingredient | Supplier | Cat # | CDD22 | CDD32 (FlyAA) | CDD33 | CDD-MM1 | CDD-MM2 | CDD-MM3 | CDD40 | CDD41 | CDD48 / HoldFast |
|---|---|---|---|---|---|---|---|---|---|---|---|
| **Amino acid mixture** | | | | | | | | | | | |
| Ultrapure Water (litres) | Millipore | Milli-Q | 0.75 | 0.75 | 0.75 | 0.75 | 0.75 | 0.75 | 0.75 | 0.75 | 0.75 |
| Sucrose | Fisher Scientific | S/8600/68 | 10.000 | 10.000 | 10.000 | 10.000 | 10.000 | 20.000 | 10.000 | 10.000 | 20.000 |
| L-Alanine | Sigma-Aldrich | A7469 | 0.500 | 1.300 | 1.810 | 2.480 | 1.220 | 1.300 | 1.810 | 1.810 | 1.810 |
| L-Arginine | Sigma-Aldrich | A8094 | 0.800 | 1.900 | 1.170 | 0.560 | 2.260 | 1.050 | 1.170 | 1.170 | 1.170 |
| L-Aspartic Acid | Sigma-Aldrich | A7219 | 0.500 | 1.370 | 1.000 | 1.200 | 1.460 | 1.500 | 1.000 | 1.400 | 1.400 |
| L-Cysteine | Sigma-Aldrich | C7352 | 0.500 | 0.400 | 0.520 | 0.035 | 0.890 | 0.350 | 0.520 | 0.550 | 0.550 |
| L-Glutamic Acid | Sigma-Aldrich | G8415 | 5.400 | 1.780 | 2.000 | 1.770 | 1.330 | 1.900 | 2.000 | 2.000 | 2.000 |
| Glycine | Sigma-Aldrich | G8790 | 0.500 | 0.900 | 1.100 | 2.270 | 0.810 | 0.600 | 1.100 | 0.600 | 0.600 |
| L-Histidine | Sigma-Aldrich | H8000 | 1.000 | 0.760 | 0.870 | 0.710 | 0.630 | 0.800 | 0.870 | 0.870 | 0.870 |
| L-Isoleucine | Sigma-Aldrich | I7403 | 3.000 | 1.310 | 1.050 | 2.130 | 0.470 | 1.200 | 1.050 | 1.400 | 1.400 |
| L-Leucine | Sigma-Aldrich | L8912 | 2.000 | 2.370 | 2.250 | 1.410 | 3.180 | 1.950 | 2.250 | 2.250 | 2.250 |
| L-Lysine HCl | Sigma-Aldrich | L8662 | 1.900 | 1.600 | 1.350 | 1.340 | 1.470 | 1.700 | 1.350 | 1.600 | 1.600 |
| L-Methionine | Sigma-Aldrich | M5308 | 0.800 | 0.700 | 0.400 | 0.560 | 0.710 | 1.350 | 0.400 | 0.700 | 0.700 |
| L-Phenylalanine | Sigma-Aldrich | P5482 | 1.300 | 1.180 | 1.120 | 0.920 | 0.960 | 2.100 | 1.120 | 1.400 | 1.400 |
| L-Proline | Sigma-Aldrich | P5607 | 0.500 | 1.150 | 1.560 | 1.060 | 1.510 | 0.950 | 1.560 | 1.200 | 1.200 |
| L-Serine | Sigma-Aldrich | S4311 | 0.500 | 1.610 | 1.400 | 1.340 | 2.560 | 1.100 | 1.400 | 1.400 | 1.400 |
| L-Threonine | Sigma-Aldrich | T8441 | 2.000 | 1.300 | 1.330 | 1.410 | 1.390 | 1.100 | 1.330 | 1.100 | 1.100 |
| L-Tryptophan | Sigma-Aldrich | T8941 | 0.500 | 0.370 | 0.270 | 0.350 | 0.210 | 0.600 | 0.270 | 0.500 | 0.500 |
| L-Tyrosine | Sigma-Aldrich | T8566 | 0.500 | 1.100 | 0.870 | 0.500 | 1.110 | 1.800 | 0.870 | 1.200 | 1.200 |
| L-Valine | Sigma-Aldrich | V0513 | 2.800 | 1.400 | 1.250 | 1.980 | 1.100 | 1.500 | 1.250 | 1.250 | 1.250 |
| L-Glutamine | Sigma-Aldrich | G8540 | 0.000 | 1.300 | 2.430 | 1.200 | 1.170 | 1.250 | 2.430 | 1.800 | 1.800 |
| L-Asparagine | Sigma-Aldrich | A4159 | 0.000 | 1.200 | 1.250 | 1.770 | 0.560 | 0.900 | 1.250 | 0.800 | 0.800 |
| Adenosine-5'(3')-MP | Sigma-Aldrich | 01930 | 0.600 | 0.600 | 0.600 | 0.600 | 0.600 | 0.600 | 0.600 | 0.600 | 0.600 |
| Guanosine-5'(3')-MP | Sigma-Aldrich | G8377 | 0.400 | 0.400 | 0.400 | 0.400 | 0.400 | 0.400 | 0.400 | 0.400 | 0.400 |
| Uridine-5'(3')-MP | Sigma-Aldrich | U6375 | 0.400 | 0.400 | 0.400 | 0.400 | 0.400 | 0.400 | 0.400 | 0.400 | 0.400 |
| Cytidine-5'(3')-MP | Sigma-Aldrich | C1006 | 0.400 | 0.400 | 0.400 | 0.400 | 0.400 | 0.400 | 0.400 | 0.400 | 0.400 |
| NaHCO3 | Fisher Scientific | S/4240/60 | 1.000 | 1.000 | 1.000 | 1.000 | 1.000 | 1.000 | 1.000 | 1.000 | 1.000 |
| KH2PO4 | Fisher Scientific | P/4800/53 | 0.710 | 0.710 | 0.710 | 0.710 | 0.710 | 0.710 | 0.710 | 0.710 | 0.710 |
| K2HPO4 | Fisher Scientific | P/5240/53 | 3.733 | 3.733 | 3.733 | 3.733 | 3.733 | 3.733 | 3.733 | 3.733 | 3.733 |
| MgSO4.7H2O | Sigma-Aldrich | M2773 | 0.820 | 0.820 | 0.820 | 0.820 | 0.820 | 0.820 | 0.820 | 0.820 | 0.820 |
| NaCl | Sigma-Aldrich | S7653 | 0.040 | 0.040 | 0.040 | 0.040 | 0.040 | 0.040 | 0.040 | 0.040 | 0.040 |
| Fe.Na EDTA | Sigma-Aldrich | EDFS | 0.020 | 0.020 | 0.020 | 0.020 | 0.020 | 0.020 | 0.020 | 0.020 | 0.020 |
| Zn.Na EDTA | Fluka | 34553 | 0.020 | 0.020 | 0.020 | 0.020 | 0.020 | 0.020 | 0.020 | 0.020 | 0.020 |
| Mn.NA EDTA | Alfa Aesar | 40519 | 0.020 | 0.020 | 0.020 | 0.020 | 0.020 | 0.020 | 0.020 | 0.020 | 0.020 |
| Cu.Na EDTA | Sigma-Aldrich | 3668 | 0.005 | 0.005 | 0.005 | 0.005 | 0.005 | 0.005 | 0.005 | 0.005 | 0.005 |
| Low-melting Agarose | Sigma-Aldrich | A9414 | 10.000 | 10.000 | 10.000 | 10.000 | 10.000 | 13.000 | 10.000 | 10.000 | 14.000 |
| **Sterol mixture** | | | | | | | | | | | |
| beta-Sitosterol | ThermoScientific | 132720050 | 0.000 | 0.000 | 0.000 | 0.000 | 0.000 | 0.200 | 0.200 | 0.200 | 0.200 |
| Cholesterol | ThermoScientific | 110190250 | 0.200 | 0.200 | 0.200 | 0.200 | 0.200 | 0.200 | 0.200 | 0.200 | 0.200 |
| Deoxycholic acid | Sigma-Aldrich | 30960 | 0.000 | 0.000 | 0.000 | 0.000 | 0.000 | 0.400 | 0.400 | 0.400 | 0.400 |
| Nipagin/Bavistin (ml)* | Sigma-Aldrich | H3647/ 378674 | 19.640 | 19.640 | 19.640 | 19.640 | 19.640 | 19.640 | 19.640 | 19.640 | 19.640 |
| **Vitamin mixture** | | | | | | | | | | | |
| Ultrapure Water (litres) | Millipore | Milli-Q | 0.25 | 0.25 | 0.25 | 0.25 | 0.25 | 0.25 | 0.25 | 0.25 | 0.25 |
| Choline Chloride | Sigma-Aldrich | C7017 | 0.0600 | 0.0600 | 0.0600 | 0.0600 | 0.0600 | 0.2000 | 0.4000 | 0.2000 | 0.2000 |
| Ca Gluconate | ThermoScientific | 211060010 | 0.0500 | 0.0500 | 0.0500 | 0.0500 | 0.0500 | 0.0500 | 0.0500 | 0.0500 | 0.0500 |
| Thymidine | Sigma-Aldrich | T1895 | 0.2000 | 0.2000 | 0.2000 | 0.2000 | 0.2000 | 0.3000 | 0.3000 | 0.3000 | 0.3000 |
| Thiamine HCl | Sigma-Aldrich | T1270 | 0.0020 | 0.0020 | 0.0020 | 0.0020 | 0.0020 | 0.0050 | 0.0040 | 0.0050 | 0.0050 |
| Riboflavin | Sigma-Aldrich | R9504 | 0.0100 | 0.0100 | 0.0100 | 0.0100 | 0.0100 | 0.0064 | 0.0064 | 0.0064 | 0.0064 |
| Nicotinic Acid | Sigma-Aldrich | N4126 | 0.0120 | 0.0120 | 0.0120 | 0.0120 | 0.0120 | 0.1200 | 0.1200 | 0.1200 | 0.1200 |
| Ca D-Pantothenate | Sigma-Aldrich | P5155 | 0.0160 | 0.0160 | 0.0160 | 0.0160 | 0.0160 | 0.0160 | 0.0160 | 0.0160 | 0.0160 |
| Pyridoxine HCl | Sigma-Aldrich | P6280 | 0.0025 | 0.0025 | 0.0025 | 0.0025 | 0.0025 | 0.0050 | 0.0040 | 0.0050 | 0.0050 |
| Folic Acid | Sigma-Aldrich | F7876 | 0.0030 | 0.0030 | 0.0030 | 0.0030 | 0.0030 | 0.0040 | 0.0040 | 0.0040 | 0.0040 |
| DL-Carnitine HCl | ThermoScientific | 108451000 | 0.0100 | 0.0100 | 0.0100 | 0.0100 | 0.0100 | 0.0100 | 0.0100 | 0.0100 | 0.0100 |
| Biotin | Sigma-Aldrich | B4639 | 0.0002 | 0.0002 | 0.0002 | 0.0002 | 0.0002 | 0.0004 | 0.0002 | 0.0004 | 0.0004 |
| Penicillin | Sigma-Aldrich | P7794 | 0.2500 | 0.2500 | 0.2500 | 0.2500 | 0.2500 | 0.2500 | 0.2500 | 0.2500 | 0.2500 |
| Streptomycin | Sigma-Aldrich | S9137 | 0.2500 | 0.2500 | 0.2500 | 0.2500 | 0.2500 | 0.2500 | 0.2500 | 0.2500 | 0.2500 |

Amounts in g unless indicated otherwise

*Nipagin/Bavistin solution is 100g Nipagin and 0.4g Bavistin in 1 litre of ethanol
