## Supplemental Table S2 for "A *Drosophila* holidic diet optimised for growth and development"

| Amino acid* | Standard diet #1 | Standard diet #2 | Autolysed yeast #1 | Autolysed yeast #2 | Cornmeal #1 | Cornmeal #2 | Agar #1 | Agar #2 |
|---|---|---|---|---|---|---|---|---|
| Ala | 0.070 | 0.072 | 2.08 | 2.09 | 0.443 | 0.435 | 0.0183 | 0.0194 |
| Arg | 0.052 | 0.051 | 1.6 | 1.62 | 0.302 | 0.29 | 0.0073 | 0.0062 |
| Asp+Asn | 0.094 | 0.096 | 3.31 | 3.4 | 0.419 | 0.399 | 0.0436 | 0.0407 |
| Cys | 0.020 | 0.023 | 0.558 | 0.669 | 0.222 | 0.223 | 0.0162 | 0.0105 |
| Cystine | - | - | - | - | - | - | - | - |
| Glu+Gln | 0.185 | 0.196 | 5.2 | 5.29 | 1.19 | 1.17 | 0.0366 | 0.0434 |
| Gly | 0.040 | 0.042 | 1.29 | 1.31 | 0.226 | 0.222 | 0.0164 | 0.0171 |
| His | 0.037 | 0.039 | 0.957 | 0.935 | 0.227 | 0.226 | 0.0177 | 0.0177 |
| Hydroxyproline | - | - | - | - | - | - | - | - |
| Ile | 0.044 | 0.048 | 1.58 | 1.59 | 0.216 | 0.215 | 0.0206 | 0.0213 |
| Leu | 0.093 | 0.098 | 2.34 | 2.38 | 0.763 | 0.749 | 0.0264 | 0.0267 |
| Lys | 0.058 | 0.058 | 2.48 | 2.51 | 0.198 | 0.189 | 0.0154 | 0.0178 |
| Met | 0.014 | 0.020 | 0.474 | 0.501 | 0.122 | 0.116 | - | - |
| Phe | 0.046 | 0.051 | 1.46 | 1.48 | 0.319 | 0.318 | 0.0264 | 0.0277 |
| Pro | 0.062 | 0.068 | 1.23 | 1.27 | 0.58 | 0.564 | 0.0273 | 0.0296 |
| Ser | 0.520 | 0.061 | 1.53 | 1.65 | 0.299 | 0.29 | 0.0175 | 0.0219 |
| Thr | 0.054 | 0.057 | 1.75 | 1.75 | 0.229 | 0.221 | 0.0277 | 0.0259 |
| Trp | 0.012 | 0.013 | 0.474 | 0.455 | 0.0501 | 0.0477 | - | - |
| Tyr | 0.038 | 0.038 | 0.896 | 0.94 | 0.143 | 0.137 | - | - |
| Val | 0.052 | 0.052 | 1.75 | 1.75 | 0.288 | 0.285 | 0.0245 | 0.0267 |
| **Total** | 1.03 | 1.08 | 30.9 | 31.6 | 6.23 | 6.09 | 0.342 | 0.353 |

*g per 100g
